# Evolution of the sex ratio and effective number under gynodioecy and androdioecy

**DOI:** 10.1101/113605

**Authors:** Marcy K. Uyenoyama, Naoki Takebayashi

**Author notes:** Corresponding author: Marcy K. Uyenoyama Department of Biology Box 90338 Duke University Durham, NC 27708-0338 USA.

## Abstract

We address the evolution of effective number of individuals under androdioecy and gynodioecy. We analyze dynamic models of autosomal modifiers of weak effect on sex expression. In our zygote control models, the sex expressed by a zygote depends on its own genotype, while in our maternal control models, it depends on the genotype of its maternal parent. Our analysis unifies full multi-dimensional local stability analysis with the Li-Price equation, which for all its heuristic appeal, describes evolutionary change over a single generation. We define a point in the neighborhood of a fixation state from which a single-generation step indicates the asymptotic behavior of the frequency of a modifier allele initiated at an arbitrary point near the fixation state. A concept of heritability appropriate for the evolutionary modification of sex emerges from the Li-Price framework. We incorporate our theoretical analysis into our previously-developed Bayesian inference framework to develop a new method for inferring the viability of gonochores (males or females) relative to hermaphrodites. Applying this approach to microsatellite data derived from natural populations of the gynodioecious plant *Schiedea salicaria* and the androdioecious killifish *Kryptolebias marmoratus*, we find that while female and hermaphrodite *S. salicaria* appear to have similar viabilities, male *K. marmoratus* appear to survive to reproductive age at less than half the rate of hermaphrodites.

## 1 Introduction

Changes in the breeding system and the effective number of individuals induce genome-wide transformations of the context in which evolution operates. Here, we address the evolution of effective number under androdioecy and gynodioecy. This analysis seeks to unify questions regarding evolutionary stability (Maynard Smith and Price 1973) of the sex ratio, the nature of heritability as defined within the Li-Price framework (Li 1967; Price 1970), and the evolution of effective number.

We begin with a description of the empirical finding of Redelings *et al*. (2015) that provides the major motivation of this study: that a measure of effective number appears to be nearly maximal in three natural populations exhibiting partial hermaphroditism (Section 1.1). Among the possible explanations for this trend is that evolution of the sex ratio in each population has coincidentally brought effective number to near maximal values (Section 1.2). However, existing theory for the evolution of the sex ratio in gynodioecious and androdioecious populations indicates that major genes for sex expression evolve to the optimal sex ratio only under complete dominance of genes inducing the development of gonochores (males or females). Accordingly, we here undertake a full analysis of the fate of rare genes of minor effect on sex expression (Section 1.3). In addition to resolving the question of the evolutionary attractiveness and stability of the optimal sex ratio, this change in perspective provides a framework for the Bayesian estimation of the viability of gonochores relative to hermaphrodites.

### 1.1 Effective number

#### Relative effective number

Wright (1931) introduced the notion of effective number in the context of generalizing fundamental aspects of evolutionary change to populations structured by sex, fluctuations through time in numbers of individuals, or other factors. In their analysis of the concept, Ewens (1982) and Crow and Denniston (1988) showed that the various definitions give rise to different expressions for effective number in even simple models.

Let *N_H_* and *N_G_* respectively denote the number of reproductive hermaphrodites and gonochores (males or females). We refer to the probability that a pair of autosomal genes, randomly sampled from distinct reproductives in the present (offspring) generation, derive from the same reproductive in the preceding (parental) generation as the rate of parent-sharing (1*/N_P_*):

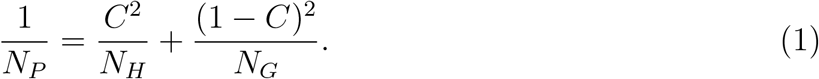

for *C* the probability that an autosomal gene randomly sampled from a reproductive in the offspring generation derives from a hermaphroditic parent. Here, we treat as equivalent the interpretation that *C* represents the collective contribution to the offspring generation from hermaphrodites in the parental generation. Descent of the gene pair from the same hermaphrodite entails that both derive from hermaphrodites (with probability *C*^2^) and from the same individual (with probability 1*/N_H_*). Similarly, the second term on the right side of (1) corresponds to the descent of the gene pair from the same gonochore. Crow and Denniston (1988) designated the inverse of the rate of parent-sharing (*N_P_*) as “inbreeding effective size.”

Redelings *et al*. (2015) defined relative effective number as the ratio of inbreeding effective size and the total effective number of reproductives (*N* = *N_G_* + *N_H_*):

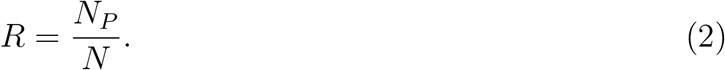

From (1), we obtain

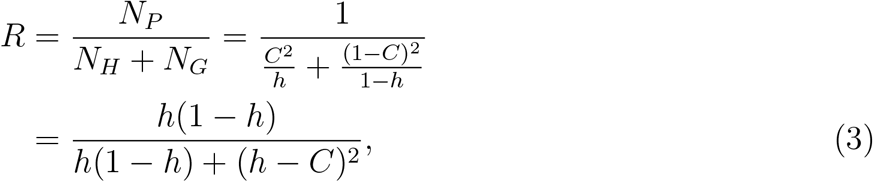

for *h* the proportion of hermaphrodites among reproductives:

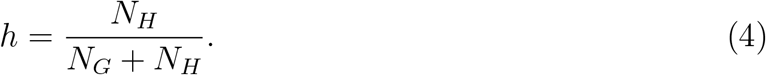

Clearly, relative effective number cannot exceed unity (*R ≤* 1), attaining unity only for

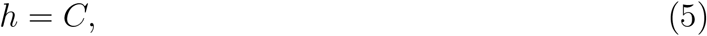

at which the proportion of hermaphrodites among reproductives (*h*) is identical to *C*, the probability that a random gene sampled from reproductives derives from a hermaphrodite in the parental generation. Both (1) and (2) differ conceptually and quantitatively from indices proposed by Laporte *et al*. (2000), who explored effective number in gynodioecious populations. That distinct concepts of effective number exist is not unexpected under even the most basic forms of population structure, including sex (Ewens 1982; Crow and Denniston 1988).

#### Empirical observations

Redelings *et al*. (2015) developed a Bayesian method for the estimation of the rate of self-fertilization in pure hermaphrodite, gynodioecious, and androdioecious populations. It provides a means of inferring all model parameters, including the determinants of relative effective number *R* (2).

Figure 1 presents posterior distributions of *R* (2) for the three data sets studied by Redelings *et al*. (2015), including those derived from two populations of the androdioecious killifish *Kryptolebias marmoratus* (Mackiewicz *et al*. 2006; Tatarenkov *et al*. 2012). An intriguing empirical observation is the near-maximization of relative effective number *R* in all three populations. A primary question motivating the present study is whether this apparent skewing reflects adaptive evolution of the sex ratio.

**Figure 1:**
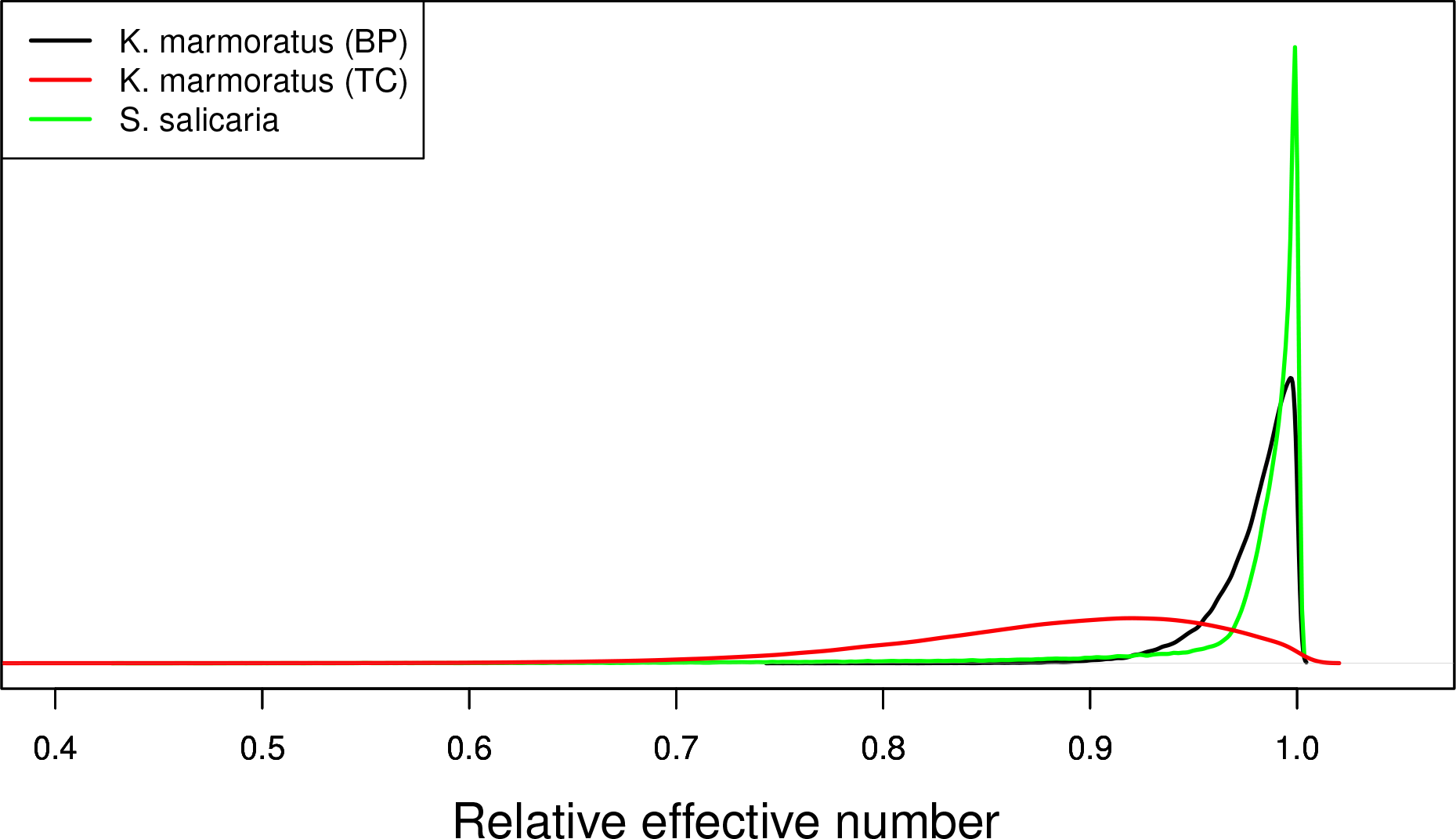
Posterior distributions of relative effective number *R* (2).

### 1.2 Evolution of the sex ratio

We address whether the near-maximal values of relative effective number (Fig. 1) reflects evolutionary pressures on the sex ratio rather than on effective number itself.

Fisher (1958) explored the evolutionary modification of the sex ratio under gonochorism, with *N_f_* females and *N_m_* males participating in reproduction. Under the assumption that reproduction is limited by the number of females, the total number of zygotes is proportional to *N_f_* and the *reproductive value* of a male relative to a female corresponds to *N_f_/N_m_*. Under gonochorism, males and females make equal collective contributions at each autosomal locus, which then implies that autosomal modifiers evolve toward equal investment in male and female offspring (Fisher 1958). Edwards (2000) provides an account of the origins of this insight.

The evolution of the sex ratio has also been addressed in the context of the marginal value of parental investment in offspring of each sex (*e.g*., Shaw and Mohler 1953; Lloyd 1975; Charnov *et al*. 1976). Increased investment in the sex with the highest marginal value affords increased transmission to the grandoffspring generation. For sexual forms corresponding to hermaphrodites and gonochores, the per capita contribution of hermaphroditic offspring to the grandoffspring generation corresponds to *C/N_H_*, reflecting the partitioning among *N_H_* reproductive hermaphrodites of the collective contribution to the gene pool by hermaphrodites (1). The marginal value of investing in hermaphroditic offspring exceeds the marginal value of investing in gonochorous offspring only if

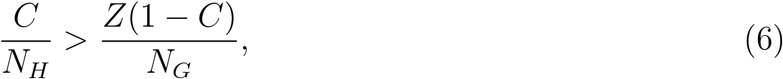

for *Z* the expected number of gonochores of reproductive age that can be produced with the investment required to produce a single hermaphrodite of reproductive age. In this context, the reproductive value of a sex is proportional to a ratio of marginal values.

An evolutionarily stable strategy (ESS, Maynard Smith and Price 1973) corresponds to an investment allocation against which no other allocation can increase when rare. Equal marginal value among mating types implies that all investment strategies give equal returns. Candidate ESS hermaphrodite proportions (*h^∗^*) among offspring at reproductive age (Adults in Table 1) correspond to points of equality between the marginal values of hermaphrodites and gonochores:

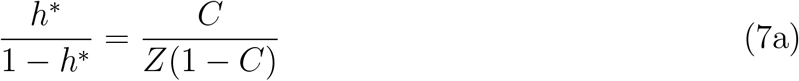

(a rearrangement of (6), using (4)). If the departure of the relative cost of a hermaphrodite (*Z*) from unity derives entirely from differential viability of gonochorous and hermaphroditic offspring between their conception and attainment of reproductive age, this candidate ESS corresponds to a sex ratio among offspring at conception (Zygotes in Table 1) of

**Table 1.**
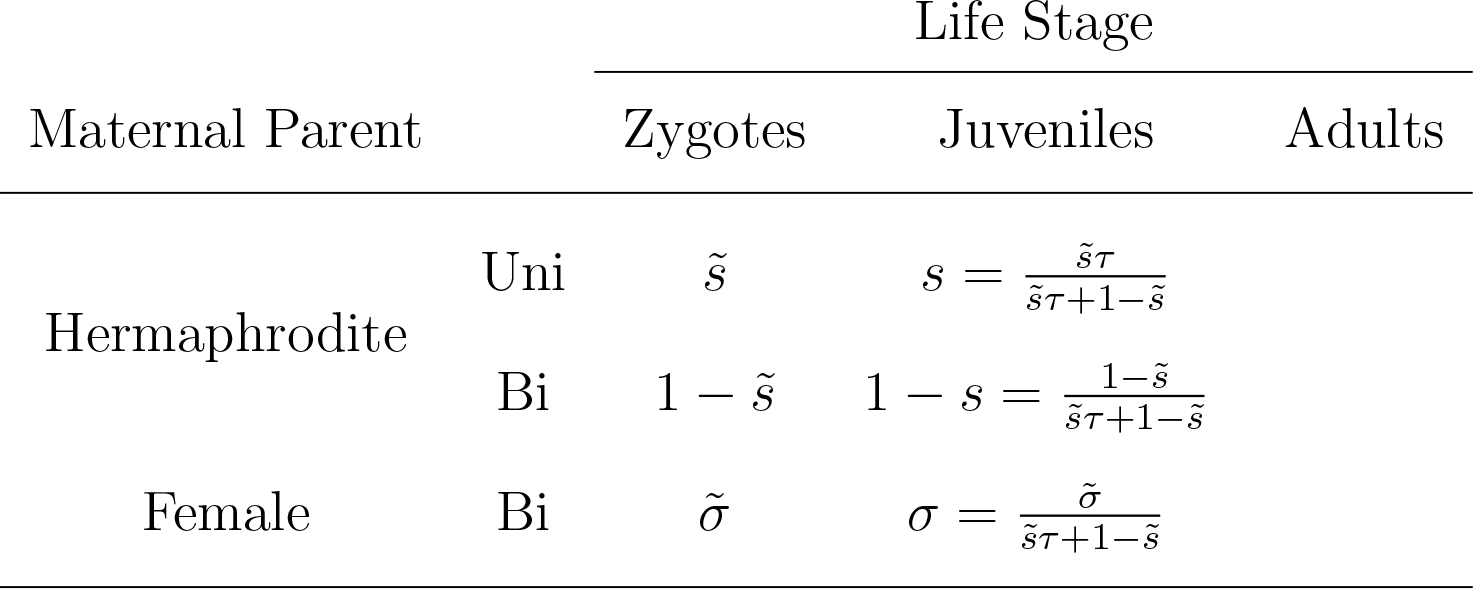
Offspring production.

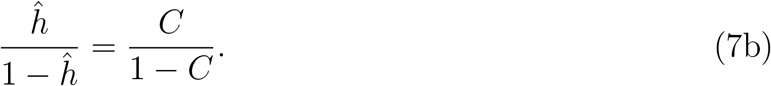

Lloyd (1975) used arguments similar to those motivating (7b) to propose ESS sex ratios under gynodioecy and androdioecy. Results of dynamic models (Ross and Weir 1975, 1976; Charlesworth and Charlesworth 1978; Wolf and Takebayashi 2004) indicate that changes in frequency of major genes for sex expression evolve to those ESS values only under complete dominance of the allele inducing gonochores (males or females), similar to a mammalian Y chromosome.

Among model systems for gynodioecy and androdioecy, evidence supporting sex determination by a dominant major gene appears to be uncommon. In their analysis of sex within broods generated by controlled crosses between females and hermaphrodites of the gynodioecious *Schiedea salicaria*, Weller and Sakai (1991) recognized two major groups of hermaphroditic pollen donors: those that generated hermaphroditic offspring almost exclusively and those that generated the female and hermaphrodites in approximately equal proportions. Weller and Sakai (1991) proposed that male sterility derives from a recessive allele at a single locus, and reported approximate agreement between the population sex ratio and the ESS proposed by Lloyd (1975). However, Ross and Weir (1975) had already shown that short-term evolution of a recessive major allele for male sterility in a gynodioecious population induces an equilibrium population sex ratio that departs from the ESS.

### 1.3 Analytical and empirical exploration

Here, we address the evolutionary modification of the sex ratio under androdioecy and gynodioecy by minor genes and its implications for effective number. We then apply this theoretical framework to empirical observations to infer *Z* (6), the relative viability of gonochores, in natural populations of the androdioecious *Kryptolebias marmoratus* and the gynodioecious *Schiedea salicaria*.

#### Evolution of the sex ratio

Among the major questions regarding the evolution of breeding systems is the nature of Darwinian fitness in this context. Reproductive success of an individual may depend not only on its own sex expression but on the sex expression of other members of the present or descendant populations. Numerous authors have explored definitions of Darwinian fitness under androdioecy and gynodioecy (Ross and Weir 1975; Lloyd 1975; Charlesworth and Charlesworth 1978). An alternative approach, and the one we have adopted here, entails modeling the genetic dynamics without appeal to an external definition of fitness (Ross and Weir 1975, 1976; Wolf and Takebayashi 2004).

While previous work has explored short-term change in the frequencies of major genes, our analysis addresses long-term change in parameter space (Eshel and Motro 1981; Taylor 1989; Christiansen 1991) by means of mutations of minor effect arising at modifier loci across the genome. A candidate hermaphrodite proportion (7b) would in fact correspond to an ESS only if any rare modifier of the sex ratio fails to increase at a geometric rate in a monomorphic population exhibiting the candidate sex ratio. Further, *ĥ* would correspond to an ESS that is locally attracting in parameter space if rare autosomal enhancers of hermaphrodite production invade a population with hermaphrodite proportion *h_c_* only if *h_c_ < ĥ* and suppressors invade only if *h_c_ > ĥ*. Such an investment allocation has been described as a continuously stable strategy (Eshel and Motro 1981) or as showing *m*-stability (Taylor 1989) or convergence stability (Christiansen 1991). We show that within the context of long-term changes in minor genes, rather than short-term changes in major genes, candidate sex ratios satisfying (7b) do in fact correspond to attracting evolutionary strategies.

Sex expression in the androdioecious killifish *Kryptolebias marmoratus* reflects epigenetic regulation of genes throughout the genome in response to temperature and other variables. Ellison *et al*. (2016) studied methylation patterns in brain tissue isolated from fish derived from eggs incubated at controlled temperatures, demonstrating significant genome wide differences due to multiple factors, including sex, temperature, and laboratory strain. Loci showing responses in methylation levels included candidates for major sex determination genes and also amplified fragment length polymorphism regions across the genome. These observations are consistent with the conceptual framework of our models, which envisions many loci with the potential to influence sex expression in response to temperature and other factors. Whether enhancement of gonochore development by a particular allele is advantageous depends on genome-wide expression patterns.

Our central theoretical result for the evolution of androdioecy and gynodioecy under both zygote and maternal control of sex expression is that the current population resists invasion of a mutation of weak effect at a modifier locus only if

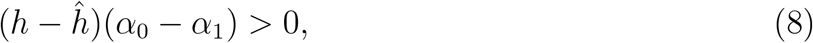

for *h* the sex expression level of the population prior to the introduction of the mutation, *ĥ* the candidate ESS sex expression level (Section 2.1), and (*α*_0_ *− α*_1_) the average effect of substitution of the new mutation inferred from the Li-Price approach extended to inbreeding (Section 2.2). This expression signifies that the population resists the introduction of a new mutation if the current sex expression level exceeds the ESS ((*h −ĥ*) *>* 0) and the average effect of the mutation would raise the sex expression level even further ((*α*_0_-*α*_1_) *>* 0), or if both conditions are reversed.

#### Estimation of relative viability

Merging our theoretical analysis with a previously-developed Bayesian inference framework (Redelings *et al*. 2015), we develop a new method for inferring the viability of gonochores (males or females) relative to hermaphrodites.

Attainment of the presumptive ESS sex ratio (7) implies maximization of relative effective number (2) only if gonochores and hermaphrodites have equal viability (*Z* = 1). The departure from unity of relative effective number provides a basis for inferring the relative viability of gonochores (*Z*). Under the assumption that the natural populations under study have in fact evolved to the ESS sex ratio at conception, we use the Bayesian sampler of Redelings *et al*. (2015) to obtain posterior densities for the relative viability of gonochores (*Z*) in *Kryptolebias marmoratus* and *Schiedea salicaria*. Our results suggest that *K. marmoratus* males have significantly lower viability than hermaphrodites in populations with both high (TC) and low (BP) frequencies of males.

## 2. Methods

### 2.1 Candidate ESS sex expression levels

We derive candidate ESS values under zygote and maternal control of sex expression in populations comprising *N_H_* hermaphrodites and *N_G_* gonochores (males or females). These candidate ESS levels extend those proposed by Lloyd (1975). Our full local stability analysis (Section 3) demonstrates that these candidates do in fact correspond to continuously stable strategies.

#### Life cycle

Table 1 summarizes offspring production by maternal parents through the major phases of the life cycle. In the androdioecy models, all maternal parents are hermaphroditic. In the gynodioecy models, females produce offspring at rate *σ̃* relative to hermaphrodites (*σ̃* corresponds to *σ* in Redelings *et al*. 2015). A proportion *s̃* of egg cells produced by hermaphrodites are self-fertilized (uniparental) and all egg cells produced by females are outcrossed (biparental). Inbreeding depression occurs between the zygote and juvenile stages, with uniparental offspring (“Uni”) surviving to the juvenile stage at rate *τ* relative to biparental offspring (“Bi”). Under a rescaling at the juvenile stage, a female has an average of *σ* surviving offspring relative to a hermaphrodite and a proportion *s* of the surviving offspring of a hermaphrodite are uniparental. Sex-specific viability selection occurs between the juvenile and adult stages, with gonochores (males or females) surviving to reproductive age at rate *Z* relative to hermaphrodites, irrespective of whether they are uniparental or biparental.

Our full dynamical models depict evolving autosomal modifiers of sex expression. In contrast, derivation of the ESS values assumes the absence of heritable variation in sex expression: for example, upon the fixation of a modifier allele that induces the ESS sex ratio. Under this assumption, offspring sex (gonochore or hermaphrodite) is independent of parental sex and independent of the level of inbreeding. Accordingly, the relative proportions of uniparental and biparental offspring (*s* and *σ* in Table 1) are identical at the juvenile and adult stages and the sex ratio among zygotes is identical to the sex ratio among juveniles. Further, C, the probability that a random gene derives from a hermaphroditic parent (1), is identical for sampling from juvenile offspring (before sex-specific selection) and from adult offspring (after sex-specific selection).

#### Androdioecy

Under androdioecy (*N_G_* males and *N_H_* hermaphrodites), outcrossing entails fertilization of egg cells from the pollen cloud, to which female-sterile (male) individuals contribute at rate *ω* relative to hermaphrodites. In accordance with the laboratory experiments of Furness *et al*. (2015) on *Kryptolebias marmoratus*, our Kryptolebias model imposes the additional assumption that all biparental individuals have a male parent (*ω* = *∞*).

Under androdioecy, all egg cells derive from hermaphrodites, with a proportion *s̃* of those egg cells fertilized by self-pollen. The uniparental proportion among juveniles,

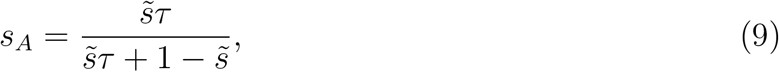

is independent of the population sex ratio.

The probability that an autosomal gene randomly sampled from juvenile offspring (Table 1) derives from a hermaphrodite in the parental generation corresponds to

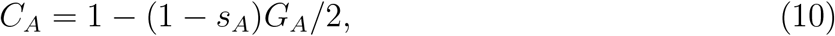

in which *G_A_* reflects the relative contribution of males of the parental generation to the pollen pool:

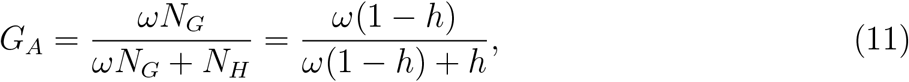

for (1 *− h*) the frequency of males among reproductives in the parental generation (4). In the Kryptolebias model, in which all biparental offspring have a male parent (*G_A_* = 1, *ω* = ∞), the collective contribution of hermaphrodites reduces to

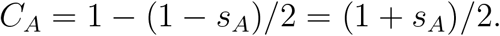

As indicated in our exposition of the life cycle (Table 1), the absence of heritable genetic variation for sex expression (*e.g*., at a genetically monomorphic ESS state) implies that the uniparental proportion *s_A_* is identical at the juvenile and adult stages. At such an ESS state, *C_A_* corresponds to the probability that a random autosomal gene derives from a hermaphrodite in the preceding generation, whether sampled at the juvenile or the adult stage.

Candidate ESS sex ratios at reproductive age (7) satisfy

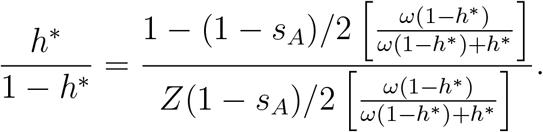

Solving, we obtain candidates for the unbeatable sex ratio at reproduction under androdioecy:

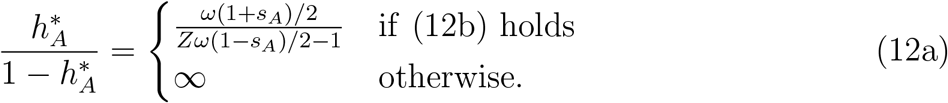

Maintenance of androdioecy (0 *< 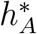 <* 1) requires that the expected contribution of a juvenile male to the subsequent generation exceed that of a juvenile hermaphrodite by at least twofold:

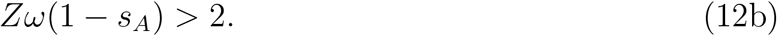

This condition becomes more stringent as the rate of outcrossing (1 − *s_A_*) or the relative viability of males (*Z*) decline. If (12b) fails, the sole candidate ESS corresponds to pure hermaphroditism (*h_A_* = 1).

At the juvenile (rather than adult) stage, the candidate ESS (12a) corresponds to a sex ratio of

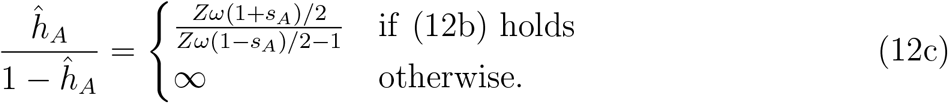

indicating that the composite parameter *Zω* represents the net effects on the ESS of differential viability and pollen success of males. Appendix A describes conditions under which the unbeatable sex ratio proposed by Lloyd (1975) corresponds to our non-zero ESS candidate (12).

#### Gynodioecy

Under gynodioecy (*N_G_* females and *N_H_* hermaphrodites), females set seeds at rate *σ̃* relative to hermaphrodites (Table 1). An autosomal gene randomly sampled from a juvenile offspring derives from a hermaphrodite parent with probability

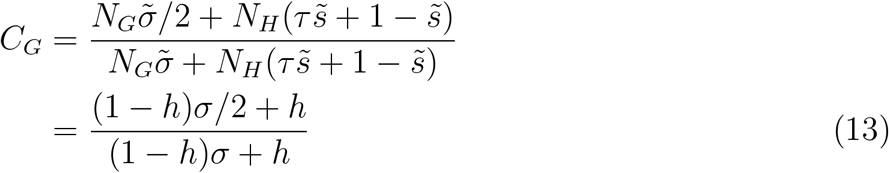

for *h* the proportion of hermaphrodites among parents in the preceding generation (4) and *σ* the scaled seed fertility of females (Table 1). This expression also corresponds to

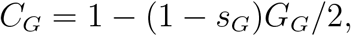

for the uniparental proportion among juveniles given by

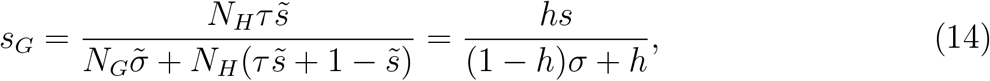

and the proportion of biparental offspring that have a female parent by

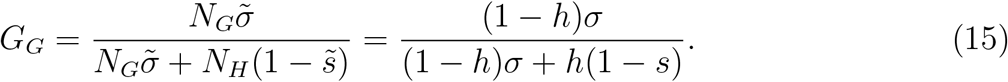

In contrast with androdioecy (9), the uniparental fraction *s_G_* (14) depends on the population sex ratio. Once again, at a monomorphic ESS (absence of heritable genetic variation for sex expression), *C_G_* provides the probability that a random autosomal gene derives from a hermaphrodite in the preceding generation, whether sampled at the juvenile or the adult stage.

From (7) and (13), the candidate ESS at reproductive age corresponds to

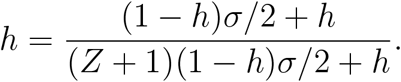

Solving, we obtain candidates for the unbeatable sex ratio under gynodioecy:

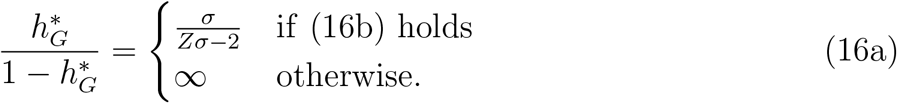

Maintenance of gynodioecy (0 *<* 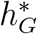*<*1) requires that the expected number of offspring produced by a juvenile female exceed that of a juvenile hermaphrodite by at least twofold:

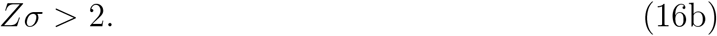

More intense inbreeding depression (smaller *τ*) and higher female viability or fertility (larger *Z* or σ ̃) tend to promote gynodioecy. For cases in which (16b) fails, the sole candidate ESS corresponds to pure hermaphroditism 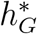 = 1).

At the juvenile stage (Table 1), the candidate ESS (16a) corresponds to

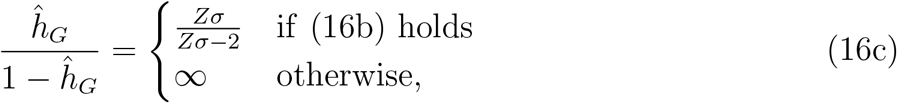

with composite parameter *Zσ* comprising the net effects on the ESS of differential viability and seed set of females. As in the androdioecy case (12), our non-zero ESS candidate (16) corresponds to the unbeatable sex ratio proposed by Lloyd (1975) under the conditions described in Appendix A.

### 2.2 Li-Price framework

Li (1967) and Price (1970) expressed the one-generation change in the frequency of an allele as a covariance between fitness and the frequency of the allele across genotypes. Here, we extend this framework to the evolution of effective number and sex ratio in inbred populations.

Table 2 presents measures associated with genotypes at a biallelic autosomal locus. In the population, genotypes *AA*, *Aa*, and *aa* occur in frequencies to *u*_0_, *u*_1_, and *u*_2_ (Σ_*i*_ u_*i*_ = 1). The locus may influence the expression of a trait, with genotype *i* associated with trait deviation (*P_i_ − P*), in which the average value of the trait corresponds to

**Table 2.** Phenotypic and genetic values.

|  | Genotypes |  |  |
| --- | --- | --- | --- |
| | $AA$ | $Aa$ | $aa$ |
| Frequency | $u_0$ | $u_1$ | $u_2$ |
| Trait deviation | $P_0 - \bar{P}$ | $P_1 - \bar{P}$ | $P_2 - \bar{P}$ |
| Additive genotypic value | $2\alpha_0$ | $\alpha_0 + \alpha_1$ | $2\alpha_1$ |
| Fitness deviation | $T(u'_0 - u_0)/u_0$ | $T(u'_1 - u_1)/u_1$ | $T(u'_2 - u_2)/u_2$ |

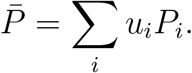

Price (1970) defined the fitness of genotype *i* as proportional to the number of gametes transmitted to the offspring generation. In panmictic populations, in which genotypic frequencies at the point of zygote formation conform to Hardy-Weinberg proportions, this definition of fitness corresponds to the expected rate of survival to reproduction, as assumed by Li (1967). Because fitness in the present context may include various components, we here define the fitness of genotype *i* as the ratio of numbers of individuals of genotype *i* at the same point in the life cycle in consecutive generations:

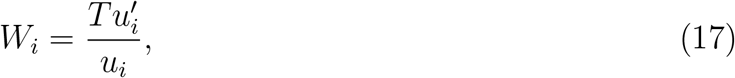

for the prime representing the next generation forward in time. Here, *T* converts the ratio of frequencies (*u^’^/u*) to the ratio of numbers of individuals; in fully-specified genetic models (Section 2.3), *T* denotes the normalizer that ensures that gene and genotypic frequencies sum to unity. Denniston (1978) observed that (17) departs from more conventional notions of fitness: high genotypic fitness reflects high production *of* the genotype rather than *by* the genotype. Under this definition, fitness is virtually always frequency-dependent: even for the most basic model of constant viability selection, (17) ceases to change only at equilibria 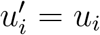

To genotypes *AA*, *Aa*, and *aa*, we associate additive genotypic values 2*α*_0_, *α*_0_ + *α*_1_, and 2*α*_1_. Much previous work, designed for panmictic populations, has defined additive genotypic value as the frequency of allele *A* in a genotype (Li 1967; Price 1970). Here, we use the definition of Fisher (1941), under which the additive effects *α_i_* are obtained by minimizing the mean squared deviation (MSD) of the phenotype from the additive genotypic value across genotypes:

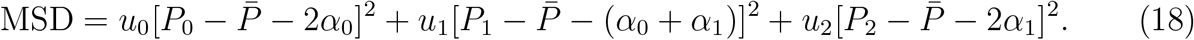

For general systems of mating, the *average effect* of substitution (Fisher 1941), the expected effect on the trait of substituting allele *A* for allele *a*, corresponds to

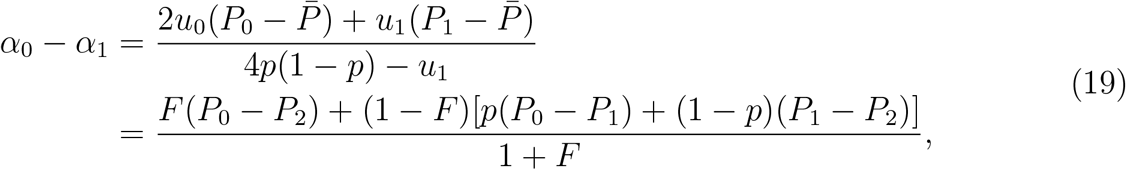

for *p* representing the frequency of allele *A* (*p* = *u*_0_ + *u*_1_*/*2) and *F* the fixation index (Wright 1933). In the additive case, in which

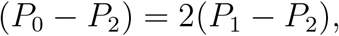

the average effect reduces to

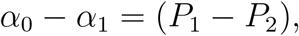

irrespective of the magnitude of *F*.

Using the definitions summarized in Table 2, we obtain the covariance across genotypes between fitness *W* (17) and additive genotypic value *G_α_* with respect to the trait:

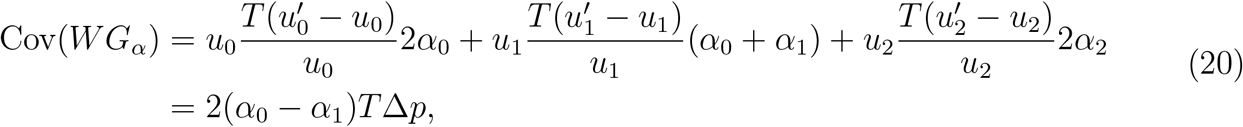

in which Δ*p* represents the change in frequency of allele *A* over a single generation. This expression indicates that the frequency of the *A* allele increases (Δ*p >* 0) if either (1) its average effect of substitution on the trait is positive ((*α*_0_ *− α*_1_) *>* 0) and the trait is positively correlated with fitness (Cov(*W G_α_*) *>* 0) or (2) its average effect of substitution on the trait is negative ((*α*_0_ *− α*_1_) *<* 0) and the trait is negatively correlated with fitness (Cov(*W G_α_*) *<* 0). To address Fisher’s (1930) fundamental theorem of natural selection, Li (1967) and Price (1970, 1971) assigned the trait of interest as fitness itself, in which case the covariance Cov(*W G_α_*) reduces to the additive variance in fitness.

For all its heuristic appeal, the Li-Price equation (20) provides a one-dimensional description of evolutionary change across a single generation. In the present context, the trait of interest corresponds to the long-term evolution of sex expression in a multi-dimensional state space. Unless sex expression is uncorrelated with fitness (Cov(*W G_α_*) = 0) or the focal modifier locus has no additive variance with respect to this trait ((*α*_0_ -*α*_1_) = 0), natural selection will induce genetic change. Because both the average effect of substitution (19) and the covariance Cov(*W G_α_*) depend on genotypic frequencies in the general case, the relationship between the one-generation description provided by (20) and the outcome of the evolutionary process needs clarification.

Key to the application of the Li-Price framework to the evolution of sex expression is the elucidation of the component of the population to which the genotypic frequencies (*u_i_*) in Table 2 correspond. In the present context, populations may comprise both gonochores and hermaphrodites, and sex expression in a zygote depends on either its own genotype or the genotype of its maternal parent. This genotypic distribution (*u_i_*) defines both the average effect of substitution (19) and heritability for the evolutionary process under study.

### 2.3 Dynamic models of sex ratio evolution

We address two genetic mechanisms for the determination of sex expression. In the zygote control models, zygotes of genotypes *AA*, *Aa*, and *aa* respectively develop into hermaphrodites at rates *h*_0_, *h*_1_, and *h*_2_ (0 *≤ h_i_ ≤* 1, *i* = 0, 1, 2), with the remaining zygotes developing into gonochores. In the maternal control models, it is the genotype of the maternal parent of the zygotes that determines sex expression rates.

Hermaphrodites set a proportion s̃ of seeds by self-fertilization. Uniparental offspring survive to reproduction at rate *τ* relative to biparental offspring, with this differential survival occurring immediately upon zygote formation, even before sex expression.

#### 2.3.1 Zygotic control of sex expression

The juvenile stage of the life cycle (Table 1) follows the expression of inbreeding depression, in which uniparental and biparental offspring may have different rates of survival, but precedes sex expression and sex-specific selection in the surviving offspring. At the end of the juvenile phase, genotypes *AA*, *Aa*, and *aa* occur in proportions *z*_0_, *z*_1_, and *z*_2_ (*z*_0_ + *z*_1_ + *z*_2_ = 1).

#### Androdioecy

In the next generation forward in time, genotypic frequencies correspond to

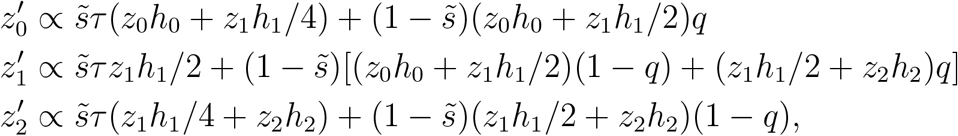

for *q* denoting the frequency of the *A* allele in the pollen pool:

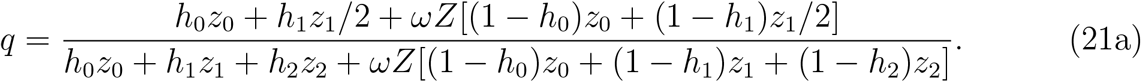

These expressions imply

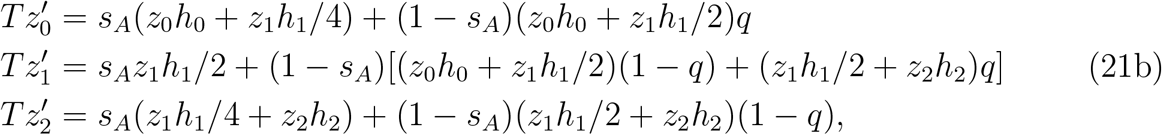

for *s_A_* given in (9) and the normalizer by

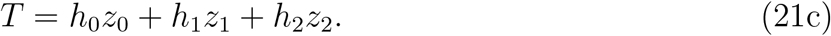

In the absence of selection on the modifier locus (*h*_0_ = *h*_1_ = *h*_2_), recursion system (21) indicates that allele frequency in seeds and pollen (*z*_0_ + *z*_1_*/*2 = *q*) remains at its initial value, with asymptotic convergence at rate *s_A_/*2 of the frequency of heterozygotes (*z*_1_) to

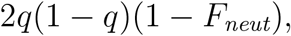

for *F_neut_* the fixation index (Wright 1933):

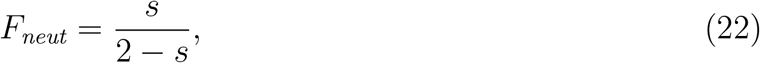

with *s_A_* substituted for *s*.

#### Gynodioecy

Genotypic frequencies in the next generation forward in time correspond to

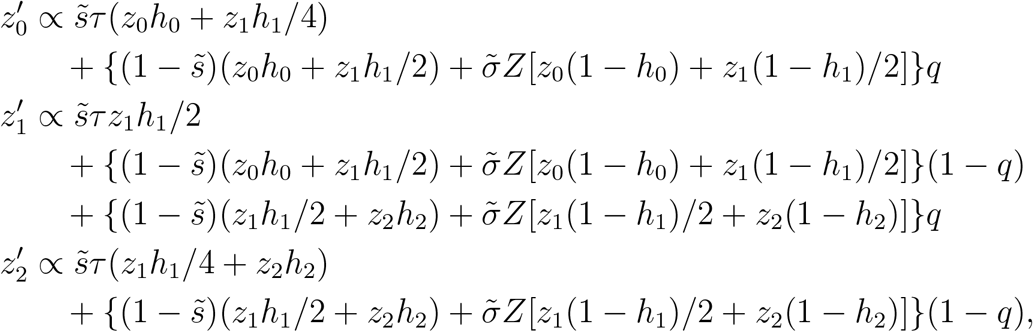

in which *q* represents the frequency of the *A* allele in the pollen pool (which derives entirely from hermaphrodites),

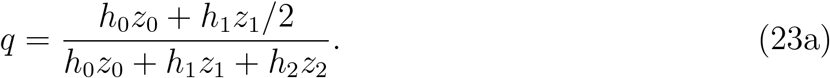

After division by (s̃*τ* + 1 – s̃), we obtain

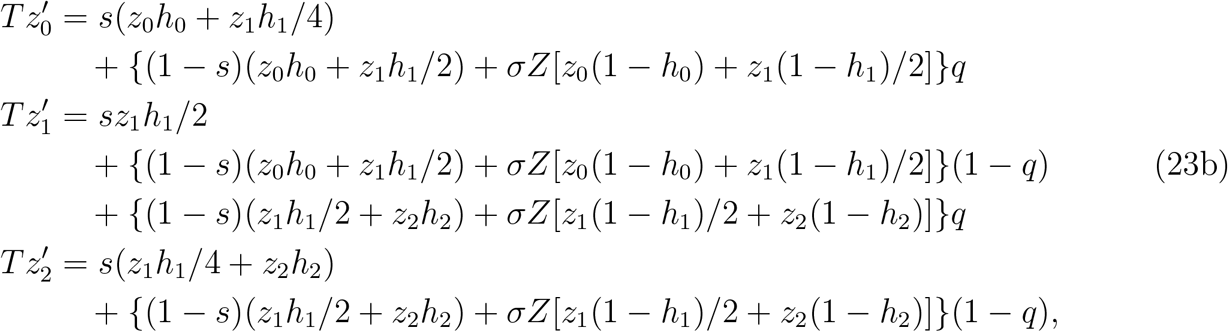

for the normalizer corresponding to

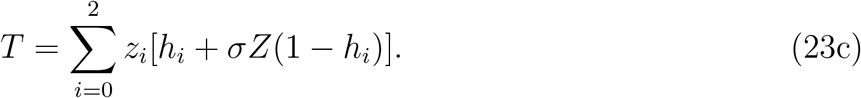

In the absence of selection on the modifier locus (*h*_0_ = *h*_1_ = *h*_2_ = *h*), allele frequency in seeds and pollen (*z*_0_ + *z*_1_*/*2 = *q*) remains at its initial value. Unlike the uniparental fraction *s_A_* (9) under androdioecy, *s_G_* (14) depends on the population sex ratio. The frequency of heterozygotes (*z*_1_) converges asymptotically at rate *s_G_/*2 (14) to

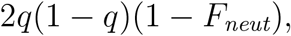

for *F_neut_* given in (22) but with *s_G_* (14) substituted for *s*. Selective neutrality at the modifier locus entails that the transformation (23) has an eigenvalue of unity (reflecting no changes in allele frequency) and an eigenvalue of *s_G_/*2 (reflecting convergence of *z*_1_ under inbreeding).

#### 2.3.2 Maternal control of sex expression

Under the maternal control model, the genotype of the maternal parent determines sex expression in zygotes. While the recursions for the zygote control models describe genotypic frequencies at the end of the juvenile stages of the life cycle, the census point for the maternal control models occurs at the end of the adult stage (Table 1), following both inbreeding depression and sex-specific selection. At this point, genotypes *AA*, *Aa*, and *aa* occur in proportions *x*_0_, *x*_1_, and *x*_2_ in hermaphrodites and *y*_0_, *y*_1_, and *y*_2_ in gonochores (*x*_0_ + *x*_1_ + *x*_2_ + *y*_0_ + *y*_1_ + *y*_2_ = 1).

#### Androdioecy

At the point of reproduction, genotypic frequencies among hermaphrodites correspond to

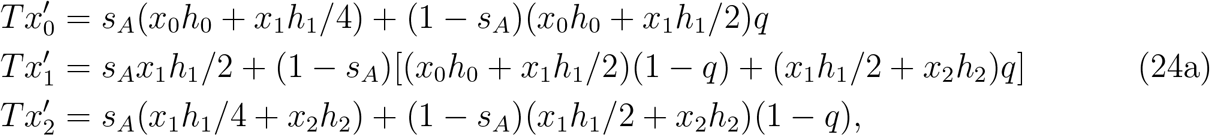

for *q* the frequency of the *A* allele in the pollen cloud,

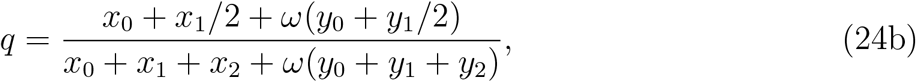

and *s_A_* given in (9). Substitution of *Z*(1 *−h_i_*) for *h_i_* in the hermaphrodite recursion 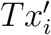 produces the male recursion 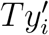, which implies the normalizer

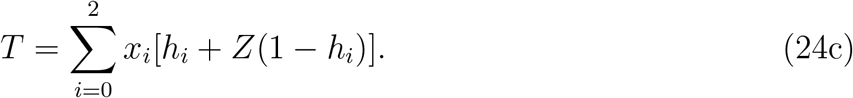

Because male genotypic frequencies (*y_i_*) affect transmission only through the pollen cloud (24b), description of the transformation requires a smaller set of variables, including *x*_0_, *x*_1_, *x*_2_, (*y*_0_ + *y*_1_*/*2), and (*y*_1_*/*2+ *y*_2_).

In the absence of selection on the modifier locus (*h*_0_ = *h*_1_ = *h*_2_ = *h*), the population ratio of hermaphrodites to males converges in a single generation to

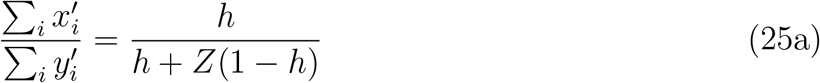

and the genotypic frequencies in hermaphrodites and males are proportional:

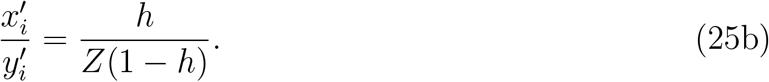

Accordingly, the frequencies of allele *A* among hermaphrodites (*x*_0_ +*x*_1_*/*2), males (*y*_0_ +*y*_1_*/*2), and pollen (*q*) converge to equality in a single generation,

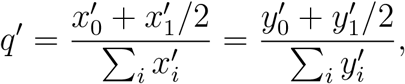

and attain their common equilibrium value in two generations,

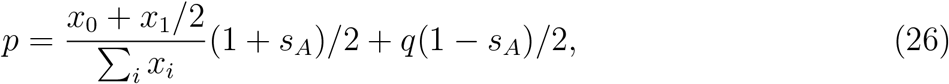

in which the uniparental proportion *s_A_* is given in (9) and *x_i_* and *q* represent the initial values of those variables. The frequency of heterozygotes converges asymptotically, at rate *s_A_/*2, to

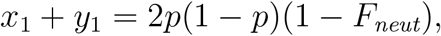

for *p* given in (26) and *F_neut_* in (22), with *s_A_* (9) substituted for *s*.

Near the state of fixation of the *a* allele, the neutral transformation has a single eigenvalue of unity (corresponding to allele frequency), a single eigenvalue of *s_A_/*2 (governing convergence of the frequency of heterozygotes to the value dictated by *F_neut_* and allele frequency), and two eigenvalues of zero (representing the near-instantaneous convergence to equality of allele frequencies in hermaphrodites, males, and pollen).

#### Gynodioecy

Genotypic frequencies in the next generation forward in time correspond to

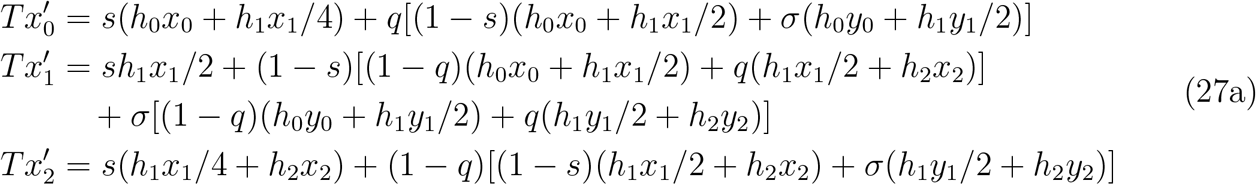

for *q* the frequency of *A* in the pollen cloud (to which hermaphrodites alone contribute):

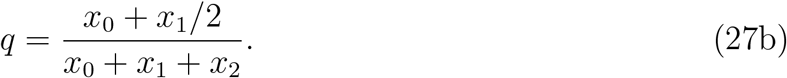

Similar to the androdioecy model, the 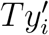 have the same form as 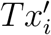 but with *h_i_* replaced by *Z*(1 *− h_i_*), which implies

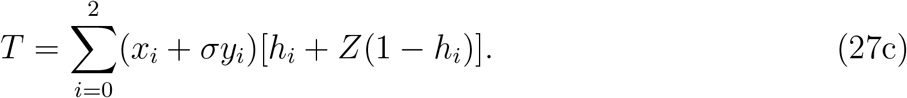

In the absence of selection on the modifier locus (*h*_0_ = *h*_1_ = *h*_2_ = *h*), the population converges in a single generation to the state (25), with the *y_i_* now representing genotypic frequencies in females. Frequencies of allele *A* among hermaphrodites, females, and pollen in the first generation correspond to

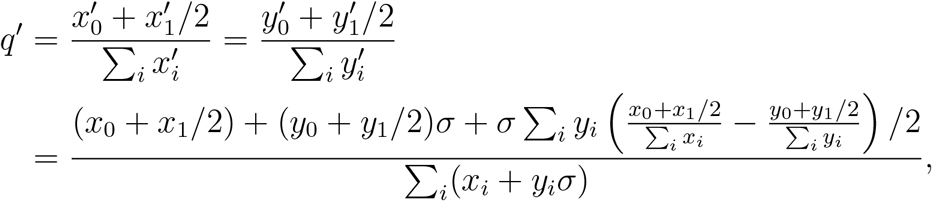

for the *x_i_* and *y_i_* representing genotypic frequencies at initialization, and attain their common equilibrium value in two generations. The frequency of heterozygotes converges asymptotically, at rate *s_G_/*2 (14), to

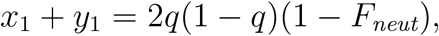

for *F_neut_* given by (22), with *s_G_* (14) substituted for *s*.

Near the state of fixation of the *a* allele, the neutral transformation has a single eigenvalue of unity (corresponding to allele frequency), a single eigenvalue of *s_G_/*2 (governing convergence of the frequency of heterozygotes to the value dictated by *F_neut_* and allele frequency), and two eigenvalues of zero (representing the convergence in two generations of allele frequencies in hermaphrodites and females to their common equilibrium value).

### 2.4 Weak selection

To explore the nature of selection on the sex ratio, we restrict most of the remaining analysis to weak selection on the modifier of the sex ratio, viewed as a perturbation from selective neutrality. Selective neutrality of the variation segregating at the focal locus entails that all genotypes induce identical hermaphrodite fractions:

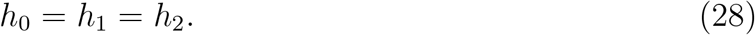

Weak selection implies that differences among genotypes,

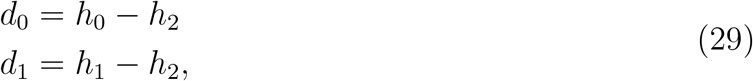

are sufficiently small to justify treating as negligible quantities of the second or higher order in the *d_i_* (*i* = 0, 1). This assumption of weak selection at the modifier locus implies no restriction on the magnitude of differences viability or fertility between inbred and outbred offspring or between the sexes.

For each of the four models under study, we determine the conditions for local stability of the fixation of the *a* allele against the introduction of the *A* allele in small frequencies. In the preceding section, we have shown that in the absence of selection on the modifier locus (28), all systems show rapid convergence to a state in which associations between genes within genotypes reflect inbreeding and associations between allele frequency and sex are absent. For each model, we enumerated the eigenvalues of the neutral transformation: a single eigenvalue of unity (representing allele frequency) and a single eigenvalue of *s/*2 (reflecting asymptotic convergence of the frequency of heterozygotes), with any additional eigenvalues corresponding to zero. Because eigenvalues are continuous in complex space (*e.g*., Serre 2010, Chapter 5), the eigenvalues of the perturbed (weak-selection) transformation depart continuously in the *d_i_* (29) from those of the neutral transformation. Accordingly, the dominant eigenvalue of the weak-selection transformation lies near unity, with the moduli of the other eigenvalues remaining strictly less than unity. Because the maternal control models have two eigenvalues of zero under neutrality, the perturbed transformation may have conjugate pairs of imaginary eigenvalues. Even so, any imaginary eigenvalues do not determine asymptotic local stability because the dominant eigenvalue of a non-negative matrix corresponds to a simple, real root of the characteristic polynomial (Gantmacher 1959). Accordingly, the dominant eigenvalue of the perturbed transformation lies near unity, with the moduli of the other eigenvalues remaining strictly less than unity. These properties of the weak-selection transformation imply that examination of the sign of the characteristic polynomial of the local stability matrix evaluated at unity is sufficient to determine local stability.

While a full local stability analysis (including, if necessary, terms of second order in the perturbations in genotype frequencies) offers a definitive determination of the fate of modifiers with weak effects on sex expression, we further undertake to elucidate the process of evolution by interpreting the results of our local stability analysis in terms of the Li-Price equation (Li 1967; Price 1970). Appendix B describes this method, which modifies an approach developed previously (Uyenoyama 1988, 1991).

## 3 Analysis

We perform local stability analyses for each of the four multidimensional models of the evolutionary modification of sex expression in androdioecious and gynodioecious populations (Section 2.3). We demonstrate that under both zygote and maternal control of sex expression, the condition for local stability corresponds to (8):

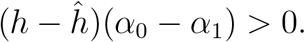

for *h* the initial sex expression level, *ĥ* the candidate ESS (Section 2.1), and (*α*_0_ *− α*_1_) the average effect of substitution (18) under the Li-Price approach extended to inbreeding.

### 3.1 Evolution of androdioecy

#### 3.1.1 Zygote control of sex expression

Under zygote control of sex expression (21), genotype *i* occurs with frequency *z_i_*, of which a proportion *h_i_* develop into hermaphrodites and the complement into males.

##### Local stability condition

A necessary condition for the exclusion of allele *A* introduced in low frequency into a population monomorphic for the *a* allele, which induces hermaphroditism at rate *h*_2_, is positivity of the characteristic polynomial of the local stability matrix evaluated at unity. Under zygote control (21), this condition corresponds to

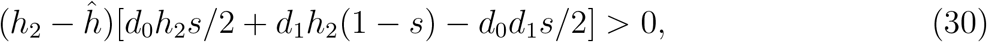

in which the uniparental proportion *s* corresponds to *s_A_* (9), *ĥ* to the ESS candidate (12c), *h*_2_ the proportion of the common *aa* genotype that develop into hermaphrodites, and the *d_i_* (29) the phenotypic deviations of genotypes bearing the rare *A* allele. Under weak selection (small *d_i_*), this condition reduces to

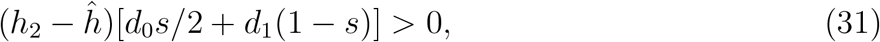

and is also sufficient for local stability. For the Kryptolebias model, in which males alone fertilize outcrossed eggs (*ω* = *∞*), we show in Appendix C that the sole condition for local stability corresponds to

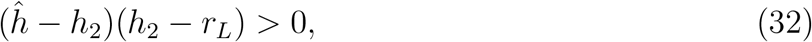

in which *r_L_* denotes the larger root of the bracketed term in (30), viewed as a quadratic in *h*_2_, under arbitrary dominance levels and intensities of selection on the modifier of sex expression (*d_i_*).

##### Average effect of substitution

A fundamental notion of heritability of sex expression is that hermaphrodites and gonochores differ in the frequencies of alleles that modify sex expression. In any generation, the difference in frequency of the *A* allele between hermaphrodites and gonochores corresponds to

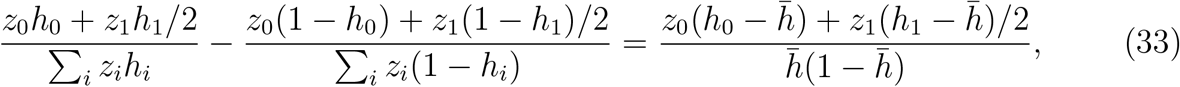

for

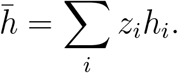

This expression corresponds to the average effect of substitution (19), with the genotypic frequencies at the point of sex expression (*z_i_*) assuming the role of the *u_i_* in Table 2.

##### New basis system

In accordance with (33), we designate as the new basis vectors near the fixation of the *a* allele (small *z*_0_ and *z*_1_)

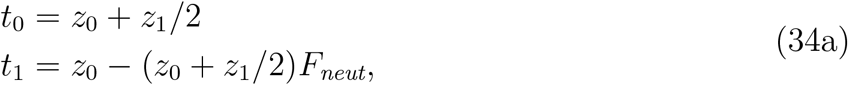

for *F_neut_* corresponding to (22) with the uniparental fraction *s_A_* (9) substituted for *s*. To the first order in the frequencies of rare genotypes, the genotypic frequencies correspond to

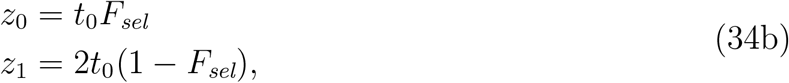

for *F_sel_* the fixation index under weak selection. From (34) we obtain

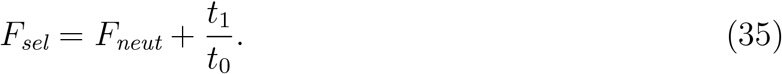

Near the fixation of the *a* allele, the average effect of substitution (19) corresponds to

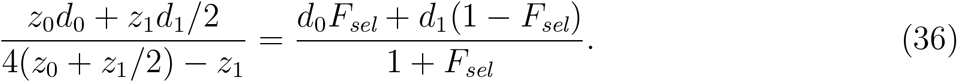

For *F_sel_* determined at the key vector (B.1) defined in Appendix B, this expression (36) for the average effect of substitution corresponds to the bracketed factor in (30).

Under weak selection (29), *t*_1_ is *O*(*d_i_*) (Appendix B), implying that the departure between *F_sel_* and *F_neut_* is also *O*(*d_i_*). To the first order in the intensity of selection on the modifier locus (*O*(*d_i_*)), the average effect of substitution (36) corresponds to

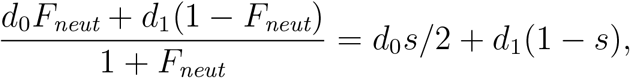

confirming that (31) corresponds to (8).

#### 3.1.2 Maternal control of sex expression

Under maternal control of sex expression (24), genotype *i* occurs with frequency *x_i_* among maternal parents, all of which are hermaphrodites, and with frequency *y_i_* among reproductive males.

##### Local stability condition

The conditions for local stability under maternal control mirror those under zygote control. The characteristic polynomial evaluated at unity is positive (necessary for local stability) only if (30) holds. Under weak selection (29), (31) provides the necessary and sufficient condition for local stability.

##### Average effect of substitution

To address heritability, we again address differences between hermaphrodites and gonochores in the frequency of a modifier of sex expression. In the next generation forward in time, the difference in frequency of the *A* allele between hermaphrodites and gonochores corresponds to

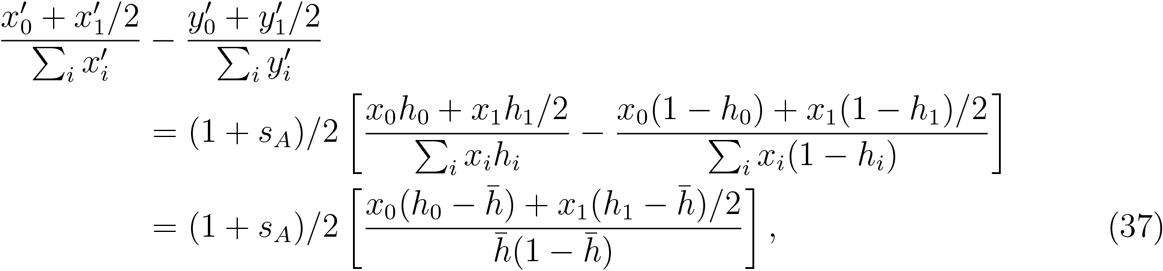

for

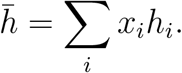

This expression suggests that the average effect of substitution corresponds to (19) with the *u_i_* replaced by

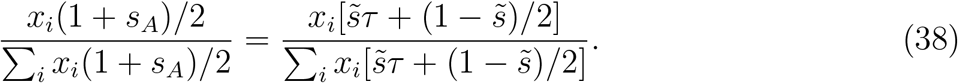

Under maternal control model of androdioecy, the maternal genotypic frequencies (*x_i_*) are weighted by the production of uniparental offspring, at rate

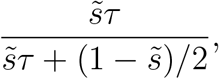

and of biparental offspring, at rate

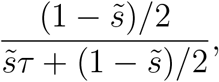

in which the 1*/*2 appears to represent the relatedness of biparental offspring to their maternal parent relative to the relatedness of uniparental offspring.

##### New basis system

We use (38) to specify the change in basis. Under androdioecy, males contribute to future generations only through pollen or sperm. In populations fixed for the *a* allele, the ratio of hermaphrodites to males at reproductive age corresponds to

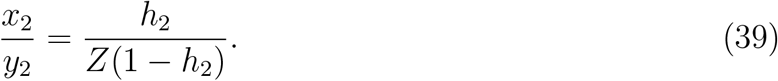

Near this fixation state, we designate as the new basis vectors

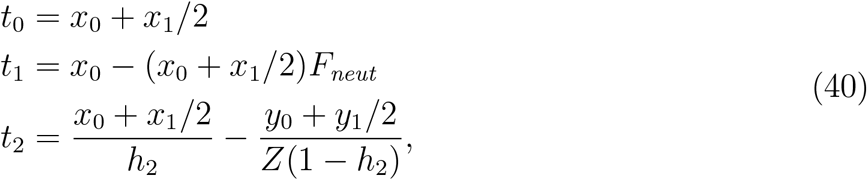

for *F_neut_* corresponding to (22) with the uniparental fraction *s_A_* (9) substituted for *s*.

At the key vector (B.1) defined in Appendix B, *t*_2_ (40), representing the difference in allele frequency between hermaphrodites and males, is proportional to the average effect of substitution (36). Also at this key vector, the fixation index under selection *F_sel_* corresponds to (35) and the average effect of substitution (36) again corresponds to the bracketed factor in (30).

Maternal and zygote control of androdioecy entail distinct definitions of the average effect of substitution (*α*_0_ *−α*_1_) (18) and the new basis system. Even so, the condition for local stability (31) again corresponds to (8).

### 3.2 Evolution of gynodioecy

#### 3.2.1 Zygote control of sex expression

For the zygote control model of gynodioecy (23), the condition for positivity of the characteristic polynomial of the local stability matrix evaluated at unity is identical to (30), with the uniparental proportion *s* now corresponding to *s_G_* (14) and *ĥ* to the ESS candidate (16c). Under weak selection (29), (31) provides the necessary condition for local stability of the fixation state.

Also identical to the expressions under zygote control of androdioecy are the average effect of substitution (33) and the definition of the new basis system (34), but with *s_G_* (14) substituted for *s* in *F_neut_* (22).

#### 3.2.2 Maternal control of sex expression

##### Local stability condition

For the maternal control model (27), the condition for local stability under weak selection corresponds to

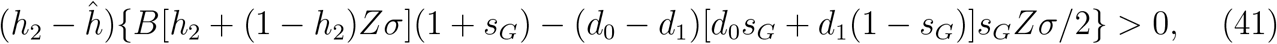

in which *s_G_* corresponds to the uniparental proportion (14), *ĥ* to the ESS candidate (16c), and *B* the bracketed factor in (30):

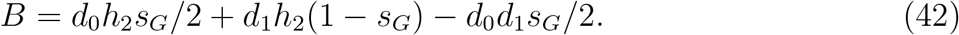

Under weak selection (29), (41) reduces to (31), which provides the necessary and sufficient condition for local stability of the fixation state.

##### Average effect of substitution

To address heritability, we return to (37). From the full system of recursions for maternal control of sex expression (27), we obtain

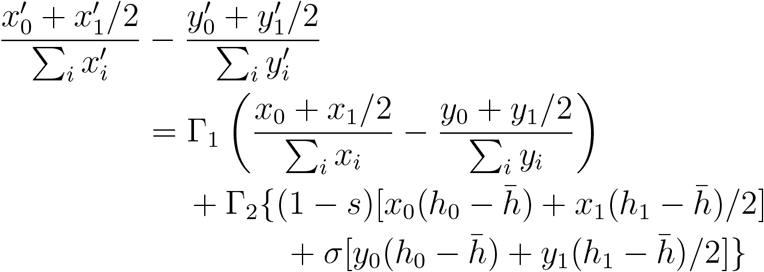

in which

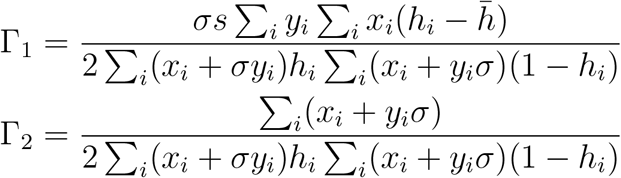

and

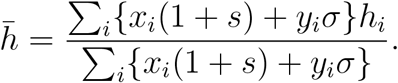

Under weak selection, for which terms of the form (*h_i_ h_j_*) are small, the difference in allele frequency between the sexes are also small, with the difference converging rapidly to

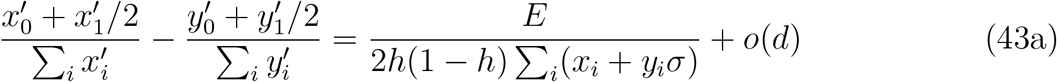

in which

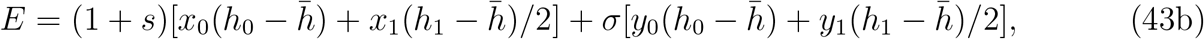

*h* represents any of the *h_i_*, and *o*(*d*) comprises quantities smaller than terms of the form (*h_i_ − h_j_*).

Expression (43b) suggests that the average effect of substitution corresponds to (19) with the *u_i_* replaced by

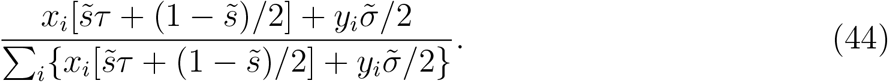

A major feature that distinguishes this gynodioecy model from the corresponding androdioecy model (37) is that gonochores (females) as well as hermaphrodites may serve as maternal parents, the individuals that control sex expression. Comparison of (38) and (44) indicates that the weighting of the contributions to the offspring generation of hermaphroditic to female maternal parents corresponds to

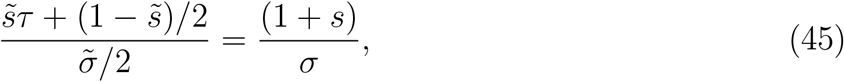

implying a twofold weighting of uniparental offspring relative to biparental offspring.

##### New basis system

In defining the new basis system, we adopt the weighted average of allele frequencies in hermaphrodites and females described in (45):

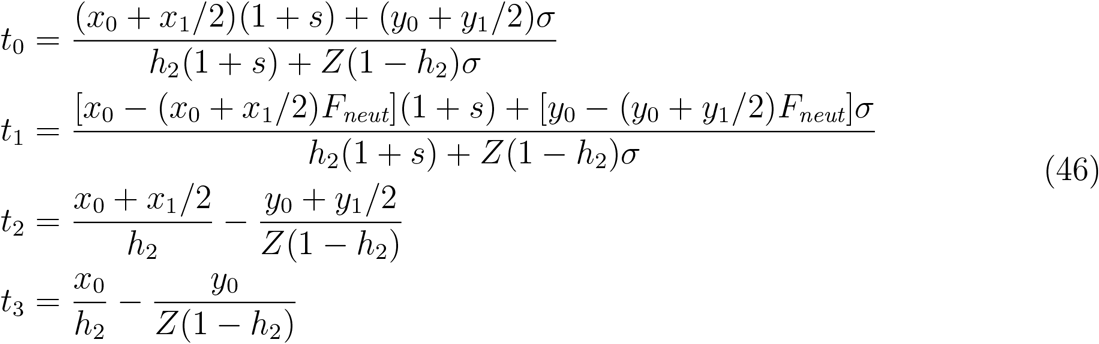

for *F_neut_* corresponding to (22) with the uniparental fraction *s_G_* (14) substituted for *s*. These expressions reflect that near the fixation state, the ratio of hermaphrodites to gonochores in the population (*x/y*) lies close to (39).

At the key vector (B.1) defined in Appendix B, both *t*_2_ (46), representing the difference in allele frequency between hermaphrodites and males, and the factor of (*h*_2_ *ĥ*) in local stability condition (41) are proportional to the average effect of substitution (43b). As in each of the other models explored, these results confirm key condition (8).

## 4 Data analysis

We use our new theoretical results to infer the viability of gonochores (males or females) relative to hermaphrodites in an androdioecious killifish and a gynodioecious Hawaiian endemic. We caution that even if a conceptual model is appropriate for a given data set, evolutionary modification over the long-term may have not yet achieved the ESS in the natural population under study.

Redelings *et al*. (2015) developed a Bayesian method for the analysis of multilocus data sampled from populations reproducing through pure hermaphroditism, androdioecy, or gynodioecy. Using an explicitly coalescent-based framework, it generates posterior distributions for the uniparental fraction (probability that a random individual is uniparental), the analogue to estimates of selfing rates generated by earlier methods (*e.g*., Ritland 2002; Enjalbert and David 2000; David *et al*. 2007).

From three microsatellite data sets derived from natural populations, Redelings *et al*. (2015) generated posterior distributions of the basic parameters of the models, including the population sex ratio among reproductives (7a). Figure 2 presents posterior distributions of the collective contribution of hermaphrodites (*C*). For the androdioecious killifish *Kryptolebias marmoratus*, the curves correspond to *C_A_* (10), and for the gynodioecious *Schiedea salicaria*, *C_G_* (13). The collective contribution of *K. marmoratus* hermaphrodites appears to be greater in the more highly inbred BP population than in the TC population. In *S. salicaria*, the (1 –*C_G_*) lies close to the population proportion of females of 12% reported by Campbell *et al*. (2010).

**Figure 2:**
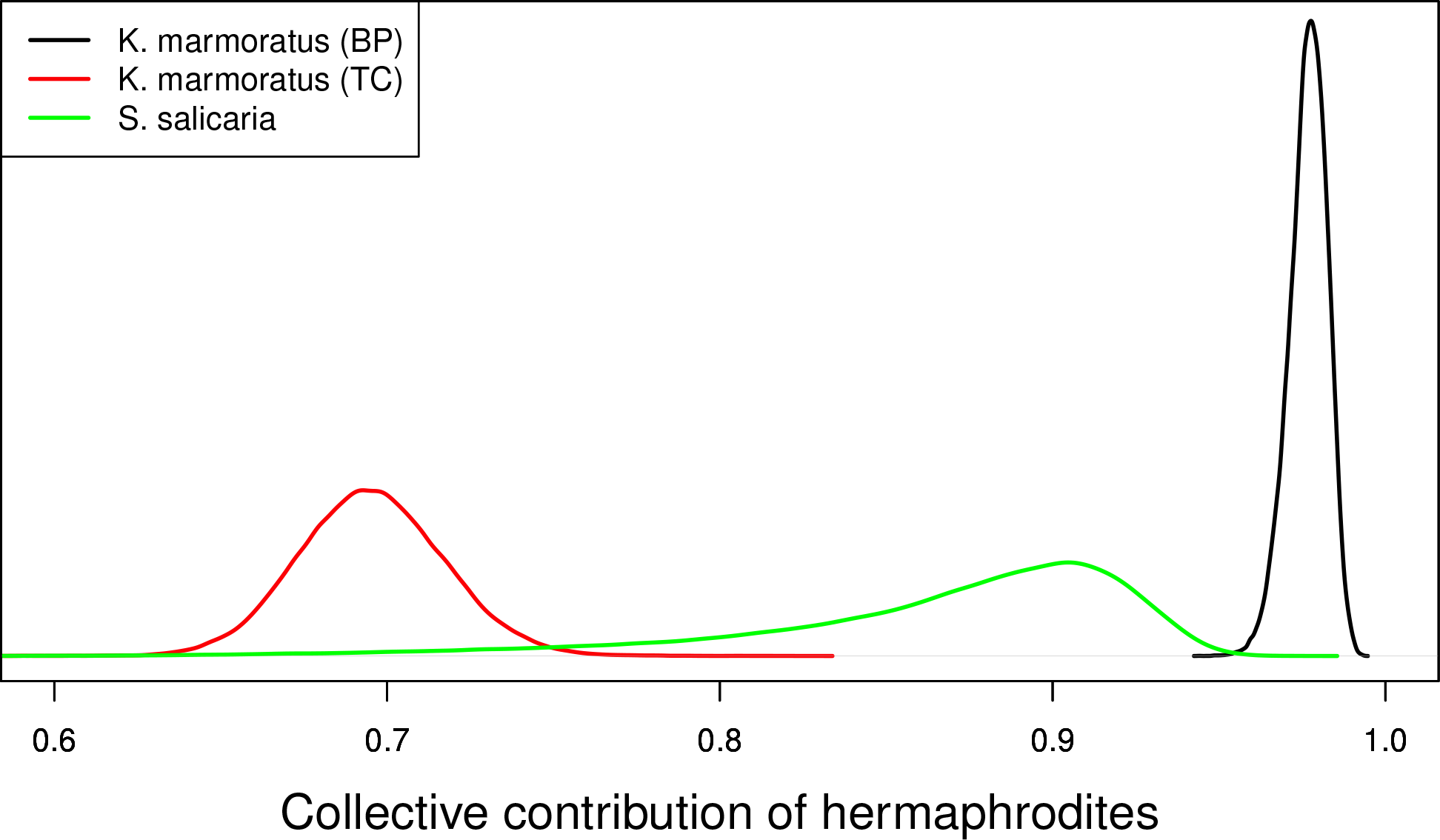
Posterior distributions of the collective contribution of hermaphrodites to the population gene pool (1) inferred from microsatellite data derived from androdioecious *Kryptolebias marmoratus* and gynodioecious *Schiedea salicaria*. For *K. marmoratus*, the curves correspond to *C_A_* (10), and for *S. salicaria*, *C_G_* (13).

From the collective contribution of hermaphrodites to the next generation (Fig. 2), we infer the sex ratio at the juvenile stage (7b). Under the assumption that the natural populations under study have converged on the attracting ESS sex ratio, we use the departure between the inferred sex ratio among adults and the expected sex ratio at conception ((7a) and (7b)) to obtain the posterior distribution of the viability of gonochores relative to hermaphrodites (*Z*).

Figure 3 presents posterior distributions of *Z* for the Schiedea and Kryptolebias populations studied. We find little evidence of a difference in viability between females and hermaphrodites in gynodioecious *S. salicaria* (median=1.08, 95% BCI=(0.34, 1.78)), in which the Bayesian Credible Interval (BCI) denotes the interval comprising the highest posterior density. In contrast, male *K. marmoratus* appear to have substantially lower viability than hermaphrodites in both the BP population (median=0.45, 95% BCI=(0.20, 0.81)) and the TC population (median=0.48, 95% BCI=(0.25, 0.77)), even though the frequency of males is several-fold higher in the TC population (0.17 versus 0.01; Turner *et al*. 1992; Tatarenkov *et al*. 2012; Mackiewicz *et al*. 2006).

**Figure 3:**
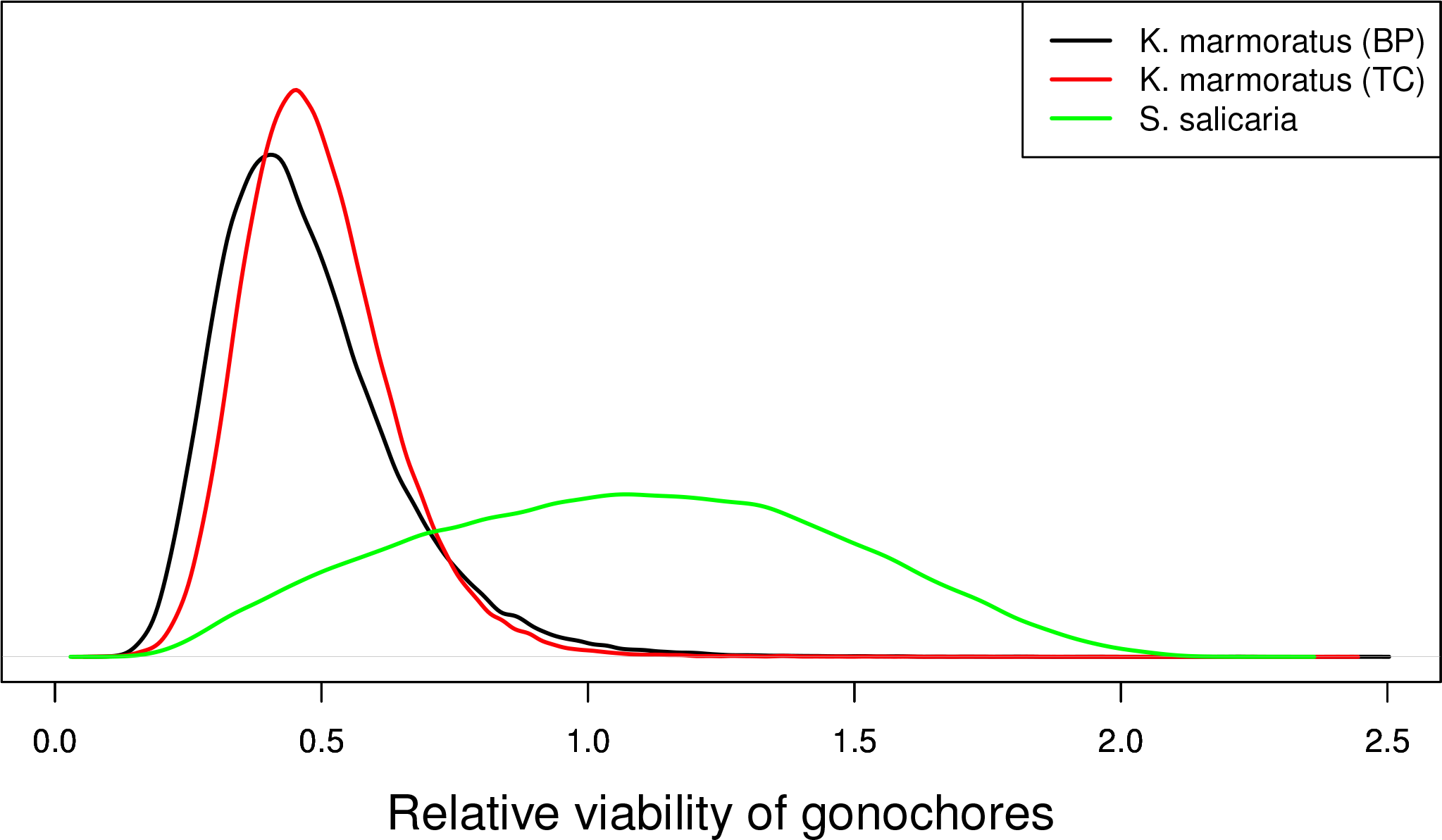
Posterior distributions of the viability of gonochores relative to hermaphrodites (*Z*).

## 5 Discussion

We have explored the evolution of androdioecy and gynodioecy under the influence of autosomal modifiers of weak effect. Our central theoretical finding (8) unifies full multi-dimensional local stability analysis with the heuristically-appealing Li-Price framework (Li 1967; Price 1970) and evolutionary stability. In addition, we have used our theoretical results to infer the viability of gonochores (males or females) relative to hermaphrodites in the gynodioecious plant *Schiedea salicaria* and the androdioecious killifish *Kryptolebias marmoratus*.

### 5.1 Relative viability of gonochores

In generalizing the Fisherian (1958) argument for the evolution of the sex ratio to androdioecy and gynodioecy, we show that *C* (1), the collective contribution of hermaphroditic parents to the offspring generation, is a central determinant of the evolutionary modification of the sex ratio. Natural selection on modifiers of weak effect promotes convergence in parameter space to the evolutionarily stable sex ratio among juveniles (Table 1) of

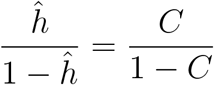

(7b), for *ĥ* the ESS proportion of hermaphrodites among juveniles and *C* corresponding to *C_A_* (10) under androdioecy and to *C_G_* (13) under gynodioecy. At reproductive age, the sex ratio corresponds to

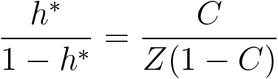

(7a), for *Z* the relative rate of survival of juvenile gonochores to adulthood and *h^∗^* the proportion of hermaphrodites after operation of sex-specific viability selection.

In the absence of sex-specific differences in rate of survival to reproductive age (*Z* = 1), relative effective number (2) is maximized (*R* = 1) at the ESS sex ratio. Here, we use the departure of relative effective number *R* from unity to infer the intensity of sex-specific differences in viability. As the Bayesian MCMC method of Redelings *et al*. (2015) permits inference of *h^∗^* and *C*, it also yields posterior distributions for *Z*, the relative viability of gonochores (Fig. 3).

The near-maximal relative effective numbers (Fig. 1) inferred for a natural population of the gynodioecious *Schiedea salicaria* (Wallace *et al*. 2011) suggests close convergence to the ESS (7b), and indeed the posterior distribution of the relative viability of gonochores (*Z*, Fig. 3) provides little evidence of differential viability between the sexes. In contrast, males of the androdioecious killifish *Kryptolebias marmoratus* (Tatarenkov *et al*. 2012) appear to have twofold lower viability than hermaphrodites (Fig. 3). Our analysis suggests a substantial reduction in male viability both in the highly inbred BP population, in which reproductively mature males are very rare (posterior median = 1%), and in the more outbred TC population, in which males are more abundant (posterior median = 17%, Redelings *et al*. 2015).

Turner *et al*. (2006) suggested that the maintenance of males in highly inbred populations of *K. marmoratus* may require “implausibly large” male fertility. Low male viability would further increase the stringency of a necessary condition (12b) for the maintenance of androdioecy under our model. However, our analysis indicates that if males alone fertilize eggs that are not self-fertilized (*ω* = *∞*, Furness *et al*. 2015), the existence of any viable biparental offspring (*s_A_ <* 1) is sufficient to favor the maintenance of males.

Turner *et al*. (2006) conducted common garden experiments to address male development in *K. marmoratus*, an emerging model system for environmental sex determination (Kelley *et al*. 2016). Lines derived from the progeny of individual hermaphrodites obtained from natural populations showed marked differences in the proportion of adult males under laboratory conditions, with fewer males in broods derived from the rare-male population. In general, laboratory-reared broods showed substantially higher frequencies of males than observed in the natural populations from which they descended. Turner *et al*. (2006) described the orange-hued mature males as “highly conspicuous.” The considerable body of work on guppies indicates that predation can generate intense selection, with various indices of crypsis responding rapidly to predator abundance under both laboratory and field conditions (Endler 1980; Reznick *et al*. 1996). The observation of an excess of *K. marmoratus* males in laboratory populations is consistent with the view that predation or other sex-specific processes operating in the wild may contribute to our inferred twofold reduction in survival to reproductive age of males relative to hermaphrodites (Fig. 3).

Our estimates of the relative viability of gonochores (*Z*) assume identity between the proportions of uniparental zygotes before (juvenile stage) and after (adult stage) the manifestation of sex-specific differences in viability (Table 1). This constraint may be violated if sex expression is heritable. Because only hermaphrodites can generate uniparental offspring, the heritability of sex may imply that hermaphroditic offspring have a higher chance both of having descended from hermaphroditic parents and of being uniparental. Consistent with this scenario is the observation (Wolff *et al*. 1988; Collin and Shykoff 2003) that gonochores appear to be more outbred than hermaphrodites in some gynodioecious species, including *Schiedea salicaria* (Weller and Sakai 2005). As a consequence, our estimates of *Z* are subject to the assumption that the population has attained the ESS level of sex expression as a genetically monomorphic state: the long-term result of filtering newly-arisen mutations of minor effect, for example.

A well-documented positive association exists between frequency of males and level of heterozygosity in natural *K. marmoratus* populations (*e.g*., Mackiewicz *et al*. 2006). Possible mechanisms include that outcrossing or high heterozygosity directly induce male development (Turner *et al*. 2006) or that the greater availability of males favors the evolution of higher outcrossing rates (Ellison *et al*. 2016). Our analysis suggests that higher outcrossing rates favor higher frequencies of males (7a), a consequence of the reduction of the collective contribution of hermaphrodites (*C*). Factors influencing the rate of outcrossing may include parasite loads, which appear to be greater in more inbred individuals (Ellison *et al*. 2011).

### 5.2 Evolution by means of major and minor genes

A considerable body of work on the evolution of gynodioecy has addressed the joint control of sex expression by major cytoplasmic and nuclear factors (reviewed by Bailey and Delph 2007; McCauley and Bailey 2009). Our analysis of autosomal modifiers does not exclude a history of cytoplasmic sex determination. For example, exclusive nuclear control may arise upon the fixation in a population of a cytoplasm that induces cytoplasmic male sterility (“cryptic CMS,” Schultz 1994; Fishman and Willis 2006). Similarly, the genetic basis of sex expression may shift from a single major locus to many loci of minor effect upon fixation at the major locus. Further, segregation of major nuclear or cytoplasmic factors does not preclude simultaneous modification of the sex ratio by nuclear factors of small effect.

#### 5.2.1 Short-term changes in frequency of major genes

Lloyd (1977) acknowledged that the frequency of a major gene inducing gonochore development converges directly to the state corresponding to the ESS (7a) only under complete dominance of the allele, under both androdioecy (Ross and Weir 1976; Wolf and Takebayashi 2004) and gynodioecy (Ross and Weir 1975). Clearly, the failure of major genes to evolve directly to the ESS under general dominance schemes is still true.

To account for the special attributes of complete dominance in existing models of short-term dynamics, we now restrict consideration to the conditions assumed by those models, including complete dominance of the gonochore allele (*e.g*., *h*_0_ = 1 and *h*_1_ = 0) and determination of zygote sex by its own genotype (zygote control model), in the absence of sex-specific selection (*Z* = 1). At the equilibrium state, let *γ* denote the proportion of offspring that have a gonochorous parent. The probability that a random autosomal gene sampled from the offspring generation at the adult stage (or juvenile stage, under *Z* = 1) derives from a gonochorous parent corresponds to

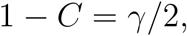

in which the half reflects the probability of sampling the allele contributed by the gonochorous parent. Half of the offspring with a gonochorous parent receive the dominant gonochore allele and themselves develop into gonochores. Accordingly, the equilibrium sex ratio in the offspring generation corresponds to

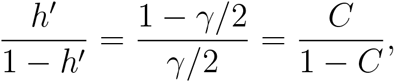

in which the half now reflects segregation of the gonochore allele and *C* denotes *C_A_* (10) under androdioecy and *C_G_* (13) under gynodioecy. This expression confirms that under complete dominance of the gonochore allele and zygote control of sex expression, the equilibrium sex ratio does indeed correspond to the ESS (7a). However, this property holds neither under other dominance schemes (*e.g*., recessive gynodioecy, Ross and Weir 1975) nor under maternal determination of zygote sex, even under complete dominance.

#### 5.2.2 Long-term modification by minor genes

Departing from earlier work on short-term changes in gene frequency space, our models accommodate general dominance among alleles of minor effect on sex expression at loci across the genome. In the androdioecious killifish *Kryptolebias marmoratus*, for example, the genome-wide epigenetic response to environmental factors documented by Ellison *et al*. (2016) is consistent with the view that loci throughout the genome may influence sex expression.

In shifting the focus to long-term changes in parameter space (Eshel and Motro 1981; Taylor 1989; Christiansen 1991), we have demonstrated that the evolutionary stability of candidate ESS values (Section 2.1) has significance beyond complete dominance of a major gene, the only domain in which short-term evolution to the ESS has been shown to occur (Ross and Weir 1975, 1976; Charlesworth and Charlesworth 1978; Wolf and Takebayashi 2004). Our results show that irrespective of the dominance of modifier genes of minor effect, long-term convergence to the ESS is expected under maternal effects on sex expression (maternal control) as well as determination of zygote sex by the zygote itself (zygote control).

Our analysis of microsatellite variation in *Schiedea salicaria* indicates near-maximal values of relative effective number (*R* near unity in Fig. 1; Redelings *et al*. 2015), supporting the view of Weller and Sakai (1991) of convergence to the ESS (7b) in the natural population studied. As noted in the Introduction (Section 1.2), their proposal of a recessive major gene for male sterility in this species is inconsistent with direct convergence of the population sex ratio to the ESS through short-term changes in the frequency of the major gene (Ross and Weir 1975). A possible reconciliation is that minor modifier loci distinct from the major gene may have induced the convergence of the population sex ratio to the evolutionarily stable strategy.

### 5.3 Local stability, evolutionary stability, and heritability

We have conducted local stability analyses for each of the four multidimensional models of the evolutionary modification of sex expression in androdioecious and gynodioecious populations (Section 2.3). Our central theoretical finding,

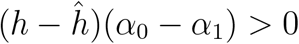

(8), implies that a new mutation that induces small effects on sex expression fails to increase when rare if the current level of hermaphroditic expression exceeds the ESS ((*h ĥ*) *>* 0) and the average effect of the mutation would raise hermaphroditic expression even further ((*α*_0_ −*α*_1_) *>* 0), or if both inequalities are reversed. We have demonstrated that under the weak-effects assumption, the conditions for local stability in multidimensional frequency space for arbitrary initial states and dominance relationships reduce to this single inequality (Section 3). Our analysis shows that the sex ratios corresponding to (7a) represent attracting evolutionarily stable strategies (ESSs) under arbitrary schemes for dominance of rare alleles at modifier loci across the genome.

#### 5.3.1 Average effect of substitution

Our central theoretical finding (8) serves to elucidate the appropriate measure of (*α*_0_ −*α*_1_), the average effect of substitution of an allele modifying sex expression in inbred populations. For general systems of mating, Fisher (1941) determined the average effect (*α*_0_−*α*_1_) by minimizing the mean squared deviation of phenotype from additive genotypic value (18). Under random mating, the average effect of substitution of a rare mutation depends only on the change in phenotype it induces in heterozygous form. Under inbreeding, determination of (*α*_0_ −*α*_1_) depends on the full array of genotypic frequencies (Section 2.2). Accordingly, demonstration of (8) for populations undergoing inbreeding requires specification of a particular genotypic array within the inherently multidimensional state space.

The heuristically-appealing Li-Price framework (Li 1967; Price 1970) provides a one-dimensional, one-generational description of the evolutionary process (20). Our unification of multidimensional local stability analysis with this framework entails specification of a key state in the full multi-dimensional space such that the change in frequency of a rare allele over a single generation starting from this state indicates its asymptotic fate (invasion or extinction) starting from arbitrary locations in a sufficiently small neighborhood of the fixation state (Appendix B). It is at this key state at which average effect (*α*_0_ −*α*_1_) in (8) is determined.

We have shown that the change in frequency of the rare allele starting from the key initial state (B.1) is proportional to the value of the characteristic polynomial, evaluated at unity, of the full transformation. Under our weak-effects assumption (29), this criterion provides necessary and sufficient conditions for local stability. However, under strong selection (introduction of genes with major effects on sex expression), the sign of the characteristic polynomial evaluated at unity is not necessarily sufficient as an indicator of local stability. In such cases, the key initial state (B.1) may become invalid or undefined (Fig. B1).

#### 5.3.2 Heritability and relatedness

The conceptual origins of the Li-Price equation (Li 1967; Price 1970) lie in Robertson’s (1966) exploration of the effects of culling, on the basis of informal criteria, on the genetic variance of a desired trait (high milk yield in dairy cattle). With respect to the evolution of mating systems, sex may influence various components of transmission of genes to future generations.

Here, the focal trait corresponds to the propensity of a zygote to develop into a gonochore or a hermaphrodite depending on the genotype at a modifier locus of its maternal parent (maternal control models) or its own genotype (zygote control models). The component of the population (zygotes or maternal parents) that influences sex expression determines the genotypic distribution (*u_i_* in Table 2) with respect to which the average effect of substitution (*α*_0_ *− α*_1_) is defined.

We adopt a notion of heritability that reflects associations between sex and allele frequency. In populations in which modifiers of sex expression segregate, the average effect of substitution (19) at a modifier locus is proportional to the difference in allele frequency between gonochores and hermaphrodites. This property holds under both zygote and maternal control of androdioecy and gynodioecy ((33), (37), (43)).

An additional question concerns the relevance of relatedness of the controlling genotype to the two sex forms (see Lloyd 1975). Under zygote control of zygote sex for both androdioecy (21) and gynodioecy (23), the average effect is defined with respect to zygote genotypic frequencies at the point of sex expression (33). In this case, the controlling entities (zygotes) are maximally related to themselves regardless of sex.

In contrast, relatedness plays a role under maternal control of sex expression. For the androdioecy model (24), hermaphrodites alone determine offspring number, with gonochores (males) serving only as pollen or sperm donors. Under our notion of heritability, the average effect is defined with respect to genotypic frequencies among maternal parents (hermaphrodites) at reproductive age (37). Uniparental offspring bear twofold higher relatedness to their maternal parents than do biparental offspring, irrespective of the sex of the offspring.

Among the unique features of maternal control of gynodioecy (27) is that gonochores (females) as well as hermaphrodites contribute to offspring number. Accordingly, the average effect of substitution depends on both sexes, with the offspring of females weighted by a factor of 1*/*2, reflecting their biparental derivation, and the biparental and uniparental offspring of hermaphrodites weighted by 1*/*2 and 1, respectively (45).

In addressing questions regarding the definition and significance of heritability and relatedness, we have permitted answers to emerge naturally from dynamic models. Our central theoretical finding (8) reconciles full multidimensional local stability analysis, ESS analysis, and the Li-Price framework. In addition, it has permitted insight into the meaning of heritability under zygote and maternal control of sex expression in inbred populations.

## Acknowledgments

We thank Bruce Turner for stimulating insights and Associate Editor Laurent Lehmann and two anonymous reviewers for their helpful comments. Public Health Service grant GM 37841 (MKU) provided partial funding for this research.

## Appendix A Lloyd’s (1975) unbeatable sex ratios

Equation (2) of Lloyd (1975) provides the unbeatable sex ratio under gynodioecy:

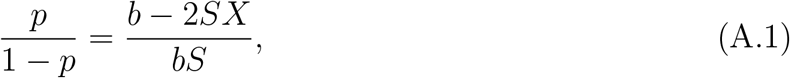

for *p* the proportion of females at reproductive age, *b* the seed set of females, *S* the viability of a hermaphrodite (described as “male”) relative to a female, and *X* the number of zygotes produced by a hermaphrodite that survive to reproduction relative to a female. In Lloyd’s notation,

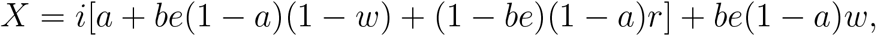

in which *i* corresponds to the relative viability of uniparental offspring (our *τ*), the bracketed quantity the proportion of seeds of hermaphrodites set by self-pollen (our *s̃*), and *be*(1−*a*)*w* the proportion of seeds of hermaphrodites set by pollen from the pollen cloud (our (1−s̃)). Substitution of

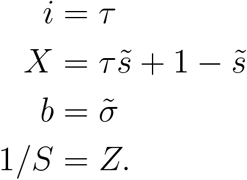

into (A.1) corresponds to our non-zero ESS candidate (16a).

In his treatment of androdioecy, Lloyd (1975, his equation (7)) proposed an unbeatable proportion of males of

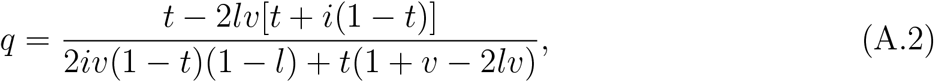

for *q* the proportion of males at reproductive age, *t* the proportion of seeds set by non-self pollen, *l* the pollen production of a hermaphrodite (described as “female”) relative to a male, *v* the rate of survival to reproduction of a hermaphrodite relative to a male, and *i* the viability of uniparental offspring relative to biparental offspring. Substitution of

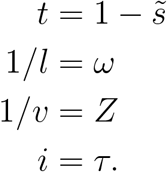

into (A.2) corresponds to our non-zero ESS candidate (12).

## Appendix B Change of basis

Here, we describe the relationship between the one-generational, one-dimensional description of evolution given by the Li-Price equation (20) and a full asymptotic, multi-dimensional local stability analysis. We describe a state of the population from which the change in allele frequency over a single generation does in fact correctly reflect the asymptotic condition for initial increase in the full multi-dimensional system under weak selection.

### B.1 Weak selection

Under selective neutrality of variation at the modifier locus, the genotypic frequencies initiated at any state comprising both alleles rapidly converge to a configuration characterized by equality between sex forms of genotypic frequencies (*x_i_* = *y_i_*) and fixation index (Wright 1933) given by (22). In the absence of differences among genotypes in sex expression (28), the multi-dimensional transformations we address (Section (2.3)) have a dominant eigenvalue of unity, reflecting preservation of allele frequency, with all remaining eigenvalues, corresponding to classical measures of disequilibrium, having moduli strictly less than unity. Weak-selection systems (29) represent perturbations in parameter space of such neutral transformations. For cases, including the maternal control model of gynodioecy (27), in which the neutral transformation has repeated eigenvalues, the perturbed transformation may have conjugate pairs of imaginary eigenvalues. Even so, any imaginary eigenvalues do not determine asymptotic local stability because the dominant eigenvalue of a non-negative matrix corresponds to a simple, real root of the characteristic polynomial (Gantmacher 1959). Because eigenvalues are continuous in complex space (*e.g*., Serre 2010, Chapter 5), the eigenvalues of the weak-selection transformation depart continuously in the *d_i_* = (*h_i_* −*h*_2_) (29) from those of the neutral transformation. In particular, the dominant eigenvalue of the local stability matrix under weak selection lies near unity, with the moduli of the other eigenvalues remaining strictly less than unity. As a consequence, examination of the value of the characteristic polynomial of the local stability matrix under weak selection is sufficient to establish local stability.

### B.2 Elucidating the Li-Price equation

To relate the Li-Price equation (20) to the full multi-dimensional local stability analysis, we introduce a change of basis from the genotypic frequencies of the rare genotypes (*AA* and *Aa*) in hermaphrodites and gonochores to allele frequency and disequilibrium measures. Here, measures of disequilibrium reflect any departures of variables other than allele frequency from their equilibrium values under the mating system in the absence of selection on the modifier locus (*h*_0_ = *h*_1_ = *h*_2_). In particular, disequilibrium corresponds to the departure of the frequency of heterozygotes (*Aa*) from the frequency associated with *F_neut_* (22) and not, in particular, from Hardy-Weinberg proportions (*F* = 0).

#### Change of basis

We determine a key vector such that the direction of change in allele frequency over a single generation starting from this vector reflects the asymptotic behavior of the system starting from an arbitrary position in the neighborhood of the fixation state. Let ***M*** denote the local stability matrix under the original basis system. Because ***M*** is a non-negative matrix, its dominant eigenvalue is non-negative and corresponds to a simple root of its characteristic polynomial (Gantmacher 1959). Under the new basis, the local stability matrix corresponds to

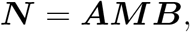

for ***A*** translating from the old basis to the new basis and ***B*** translating from the new basis to the old basis (***AB*** = ***I***). For ***z*** an arbitrary vector in the neighborhood of the fixation state,

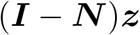

describes change over a single generation. We define key vector z̃ such that change may occur only in the first dimension (allele frequency), irrespective of the magnitude of disequilibria in other dimensions:

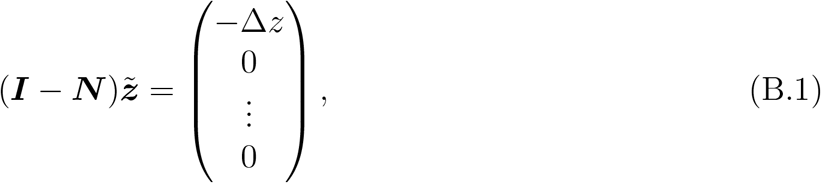

in which Δ*z* denotes the change in allele frequency over a single generation. For ***M*** and ***N** n*-dimensional matrices, z̃ is determined by the last (*n −* 1) rows of (***I** − **N***)z̃.

#### Asymptotic behavior

Here, we show that under weak selection (29), a one-generation step from key vector z̃ (B.1) indicates the asymptotic behavior of the system initiated from an arbitrary location in the neighborhood of the fixation state.

Let ***X*** represent the matrix obtained by replacing the first column of an *n*-dimensional identity matrix by z̃. Multiplication of (***I** − **N***) by ***X*** on the right produces

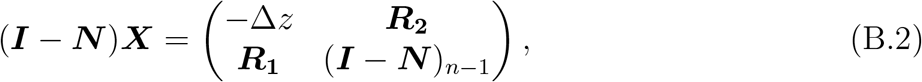

in which ***R***_1_ is an (*n −* 1)-dimensional column vector of zeros, ***R***_2_ is an (*n −* 1)-dimensional row vector with elements equal to the corresponding elements of the first row of (***I** − **N***), and (***I** −**N***)_*n−*1_ is the matrix obtained by removing the first row and column from (***I** −**N***). Taking the determinant of both sides of (B.2) produces

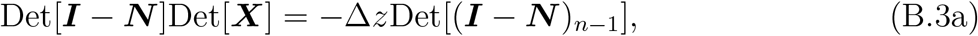

for Det[(***I** − **N***)_*n−*1_] the principal minor obtained by deleting the first row and column of (***I** − **N***).

To achieve our objective of relating the Li-Price equation (20) to a full multi-dimensional local stability analysis, we demonstrate that

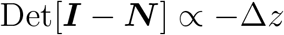

under weak selection (*d_i_* near zero). This expression implies that the direction of change over a single generation of the system initiated at z̃ (B.1) corresponds to the sign of the characteristic polynomial of the multi-dimensional stability matrix evaluated at unity. Weak selection (29) entails small differences among genotypes in sex expression (small *d_i_* = *h_i_ −h*_2_). Because Δ*z* is *O*(*d_i_*), (B.3a) implies

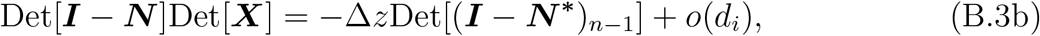

for ***N ^∗^*** the linearized transition matrix under neutrality (*d_i_* = 0). To show that

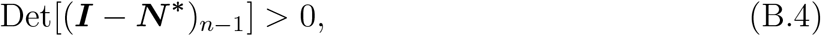

we note that under neutrality, the absence of all disequilibria implies invariant gene frequency in all models studied here (Section 2.3). Accordingly,

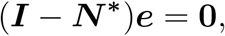

in which ***N ^∗^*** denotes the linearized transition matrix under neutrality and ***e*** the unit vector with first element equal to 1 and zeros elsewhere. This expression implies that the element in the first column and row of ***N ^∗^*** corresponds to unity. Further, that the neutral system converges to the state in which all disequilibria are absent implies that all elements in the first column of ***N ^∗^*** other than the first are zero. As a result, ***N ^∗^*** has the form

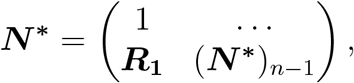

in which (***N ^∗^***)_*n−*1_ denotes the submatrix obtained by removing the first row and column from ***N ^∗^*** and ***R***_1_ is again an (*n*−1)-dimensional column vector of zeros. The characteristic polynomial of ***N ^∗^***,

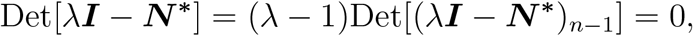

has a unit eigenvalue (corresponding to allele frequency), with the remaining eigenvalues (corresponding to disequilibria) given by the roots of Det[(*λ**I*** −***N ^∗^***)_*n−*1_]. That all eigenvalues associated with disequilibria are strictly less than unity in absolute value implies (B.4).

## Appendix C Local stability analysis of Kryptolebias model under zygote control of sex

We address the local stability of recursion system (21) near the state of fixation of the *a* allele at the modifier locus. Under Kryptolebias model, sex expression is determined by the genotype of the zygote itself and only males fertilize outcrossed eggs (*ω* = ∞ and *Z >* 0). In the absence of males prior to the introduction of genetic variation at the modifier locus (*h*_2_ = 1), eggs that are not self-fertilized fail to become zygotes. As a consequence, any allele that induces the development of males (*h*_0_ *>* 0 or *h*_1_ *>* 0) derives an enormous selective advantage from the fertilization of the proportion (1−*s_A_*) of all eggs produced. Accordingly, we restrict further consideration to cases in which the common genotype produces some males (*h*_2_ *<* 1).

We demonstrate that the sole condition for local stability corresponds to (32):

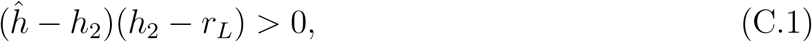

for *r_L_* the larger root of the bracketed term in (30), viewed as a quadratic in *h*_2_:

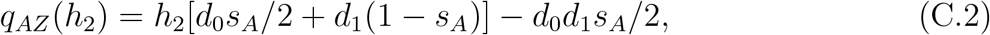

in which

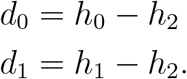

These results imply that the proposed ESS (12c) corresponds to an attracting evolutionarily stable strategy under arbitrary dominance levels and intensities of selection on the modifier of sex expression.

### C.1 Linearized recursion system

At the fixation of the *a* allele, the population comprises only *aa* individuals (*z*_2_ = 1), with normalizer

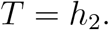

Upon the introduction of the rare alternative allele *A*, genotypes *AA* and *Aa* arise in low frequencies (*δ*_0_ and *δ*_1_). Linearization of the full recursion system (21) by ignoring terms of the second order in the *δ_i_* produces

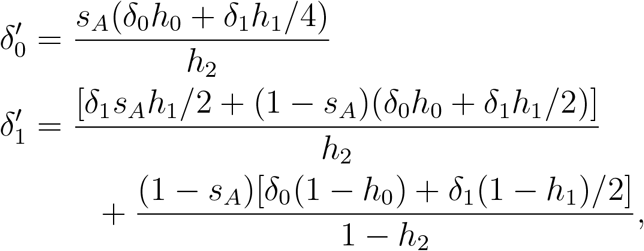

with local stability determined by the dominant eigenvalue of

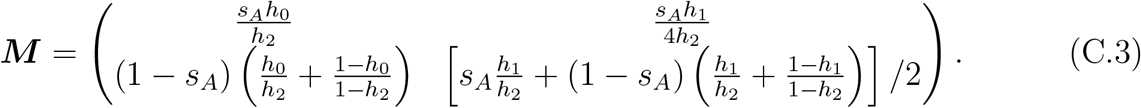

Because this matrix is non-negative, its dominant eigenvalue is real and non-negative (Gantmacher 1959, Chapter XIII). Its characteristic polynomial is proportional to

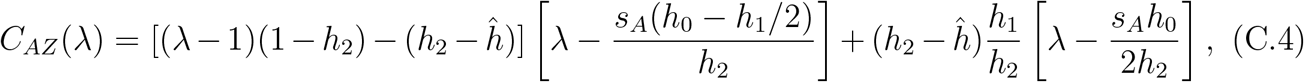

in which the proposed ESS proportion of hermaphrodites at birth corresponds to (12c), with all biparental offspring derived from male parents (*ω* = *∞*):

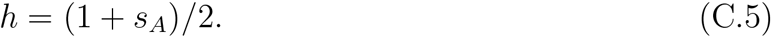

Setting the resident hermaphrodite fraction to the proposed ESS (*h*_2_ = *ĥ*), we find that *C_AZ_* (*λ*) (C.4) reduces to

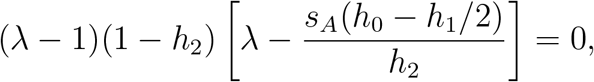

confirming a dominant eigenvalue of unity near the fixation of an allele that induces the candidate ESS, as required for an ESS. Further, we show that ESS is evolutionarily attracting: in a population fixed for an allele that specifies a sex ratio different from the ESS (*h*_2_ ≠*ĥ*), only alleles that locally bring the sex ratio closer to the ESS increase when rare (C.1).

A necessary condition for local stability is positivity of the characteristic polynomial *C_AZ_* (*λ*) (C.4) evaluated at unity. In addition, we determine the sign of *C_AZ_* (*λ*) at two values:

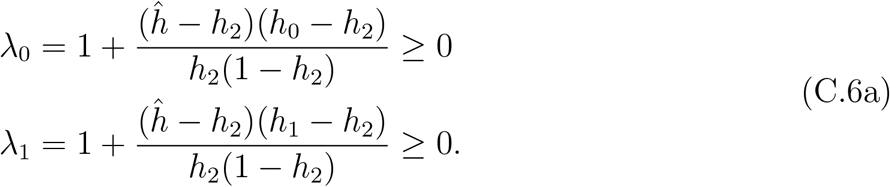

We find that *C_AZ_* (*λ*) changes sign between these values:

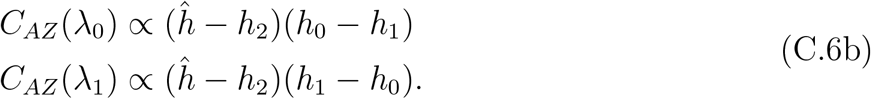

### C.2 Special cases

Under random mating (*s_A_* = 0), the ESS *ĥ* (C.5) reduces to 1*/*2 and the dominant eigenvalue of local stability matrix ***M*** (C.3) corresponds to

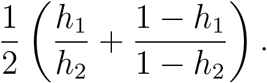

This condition implies that the fixation state is locally stable only if

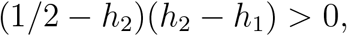

confirming both (C.1) and the classical results of Fisher (1958, Chapter VI): an equal sex ratio at birth corresponds to an attracting ESS under random mating.

Under complete selfing (*s_A_* = 1), the ESS *ĥ* (C.5) is equal to unity. Matrix ***M*** (C.3) is triangular, with the fixation state locally stable to the introduction of the *A* allele only if

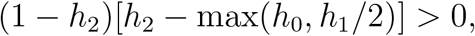

again confirming (C.1).

Under complete dominance of the rare allele (*h*_0_ = *h*_1_), characteristic polynomial (C.4) reduces to

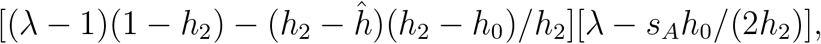

and the larger (*r_L_*) and smaller (*r_S_*) roots of (C.2) correspond to

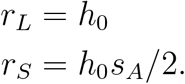

Local stability requires that both

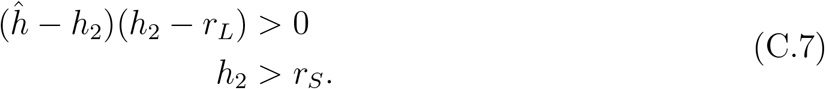

Because

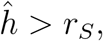

the necessary and sufficient condition for local stability under complete dominance corresponds to the first inequality in (C.7), in accordance with (C.1).

### C.3 General dominance and selection intensity

In the remainder of this section, we assume partial inbreeding (0 *< s_A_ <* 1) and *h*_0_ ≠ *h*_1_. We first demonstrate that (C.1) implies positivity of the characteristic polynomial (C.4) evaluated at unity for all *h*_0_ and *h*_1_. We then show that this necessary condition for local stability is in fact sufficient: the (non-negative) dominant eigenvalue of ***M*** (C.3) is less than unity under (C.1).

Substitution of *λ* = 1 into the characteristic polynomial (C.4) indicates

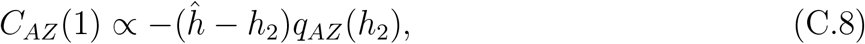

for *q_AZ_* (*h*_2_) given in (C.2). In accordance with our earlier exposition of the full recursion system (21), neutrality (*d*_0_ = *d*_1_ = 0) implies that the eigenvalue associated with allele frequency corresponds to unity, with the frequency of *Aa* heterozygotes converging to the state corresponding to *F_neut_* (22) at rate *s_A_/*2.

We now assume that *d*_0_ or *d*_1_ is non-zero (*h*_0_ ≠ *h*_2_ or *h*_1_ ≠ *h*_2_). Because only hermaphrodites produce egg cells, the existence of the population monomorphic for the *a* allele implies *h*_2_ *>* 0. If the rare allele determines complete male development (*h*_0_ = 0 or *h*_1_ = 0), then smaller root *r_S_* = 0 and

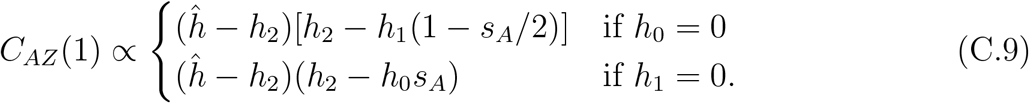

If *h*_0_ = 0 and *C_AZ_* (1) *>* 0, then (C.6) indicates that

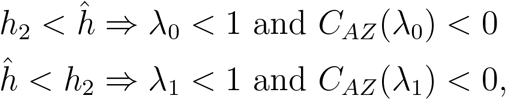

Similarly, under *h*_1_ = 0 and *C_AZ_* (1) *>* 0,

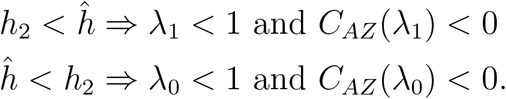

These relationships indicate the existence of a root of characteristic polynomial *C_AZ_* (*λ*) in (0, 1), which confirms (32) and (C.1): *C_AZ_* (1) *>* 0 is both necessary and sufficient for local stability under *h*_0_ = 0 or *h*_1_ = 0.

Restricting consideration to the remaining case (*h*_0_, *h*_1_, *h*_2_ *>* 0), we find that *q_AZ_* (*h*_2_) (C.2) corresponds to a quadratic in *h*_2_ with a negative leading term with

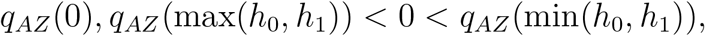

which implies that the larger (*r_L_*) and smaller (*r_S_*) roots of this quadratic lie in

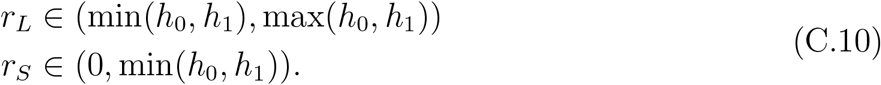

We first establish that

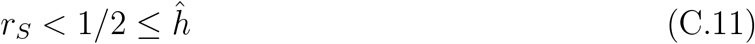

for all *h*_0_ and *h*_1_ in (0, 1]. In cases satisfying

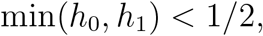

the smaller root *r_S_* (C.10) lies below 1*/*2 and consequently *ĥ*. If

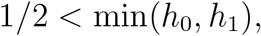

both *d*_0_ and *d*_1_ are positive for *h*_2_ = 1*/*2, which implies

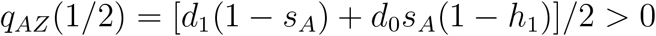

and confirms (C.11).

For small *h*_2_, satisfying

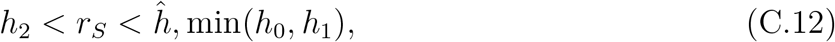

*C_AZ_* (1) *>* 0 (C.8) and both *λ*_0_ and *λ*_1_ exceed unity (C.6). That the quadratic characteristic polynomial (C.4) is negative at one of these values (*C_AZ_* (*λ*_1_) *<* 0 or *C_AZ_* (*λ*_0_) *<* 0) implies that an eigenvalue in excess of unity exists. We conclude that under (C.12), alleles that increase the proportion of hermaphrodites beyond the level specified by the resident homozygote (*h*_2_) increase when rare, confirming (C.1).

We now consider higher hermaphroditic frequencies at the fixation,

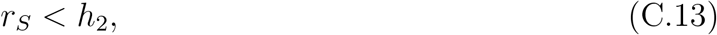

under which (C.8) indicates

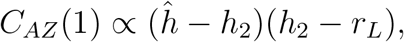

the left side of (C.1). Accordingly, (C.1) (*C_AZ_* (1) *>* 0) is a necessary condition for local stability. We now demonstrate that it is in fact sufficient for local stability under (C.13). For

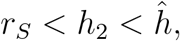

*C_AZ_* (1) *>* 0 implies

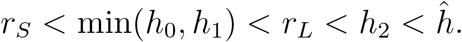

Expressions (C.6) indicate that if *h*_1_ *> h*_0_, *λ*_0_ *<* 1 and *C_AZ_* (*λ*_0_) *<* 0 If *h*_0_ *> h*_1_, *λ*_1_ *<* 1 and *C_AZ_* (*λ*_1_) *<* 0. We conclude that quadratic characteristic polynomial (C.4) is negative at a value (*λ*_0_ or *λ*_1_) less than unity, which implies that that *C_AZ_* (1) *>* 0 (C.1) is sufficient for local stability. We now restrict consideration to

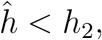

which together with *C_AZ_* (1) *>* 0 implies

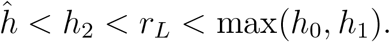

Similar to the preceding case, *h*_1_ *> h*_0_ implies *λ*_1_ *<* 1 and *C_AZ_* (*λ*_1_) *<* 0, while *h*_0_ *> h*_1_ implies *λ*_0_ *<* 1 and *C_AZ_* (*λ*_0_) *<* 0 (C.6). We again conclude that the quadratic characteristic polynomial (C.4) has a root less than unity, which implies that (C.1) is indeed necessary and sufficient for local stability.

### C.4 Limits of the Li-Price equation

Here, we illustrate that the weak-selection assumption is essential to the heuristically-appealing Li-Price equation (20) and the change of basis that relates it to a full local stability analysis. We provide an example showing that under strong selection on the modifier locus, the key vector z̃ (B.1) can become invalid and the sign of the characteristic polynomial evaluated at unity insufficient to determine the asymptotic fate of a rare allele introduced into a population monomorphic at the modifier locus.

Local stability matrix ***M*** (C.3) represents the linearized transformation with respect to a basis comprising the frequencies of rare genotypes *AA* (*δ*_0_) and *Aa* (*δ*_1_). We adopt the new basis described in Appendix B, which comprises the frequency of the rare allele (*A*) and the departure of the heterozygote frequency from the proportion expected under neutrality (*h*_0_ = *h*_1_ = *h*_2_):

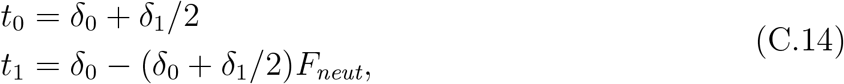

for

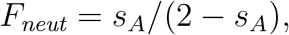

the fixation index under uniparental fraction *s_A_* (9). Matrix ***A***,

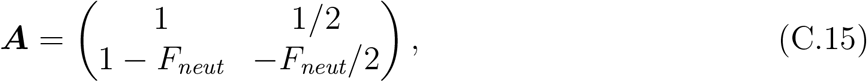

translates points from the original to the new coordinate system. In the original coordinate system, the key vector (B.1) z̃ corresponds to

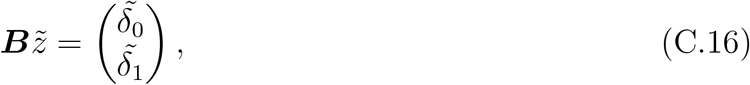

for ***B*** = ***A**^−^*^1^.

For illustrative purposes, we assume additivity in sex expression,

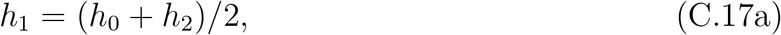

and set

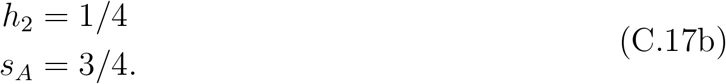

Under these assignments, the ESS *ĥ* evaluated at unity (C.8) reduces to corresponds to 7*/*8 and the characteristic polynomial

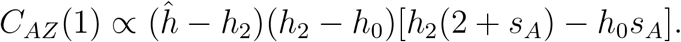

The sole condition for local stability (C.1), which reduces to

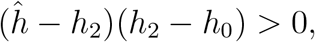

indicates that the fixation of the *a* allele resists the invasion of the rare *A* allele only for

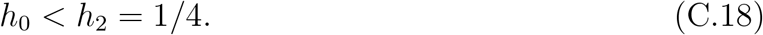

Indeed, the characteristic polynomial evaluated at unity *C_AZ_* (1) is positive in this range and changes sign at *h*_0_ = 1*/*4. However, under intense selection,

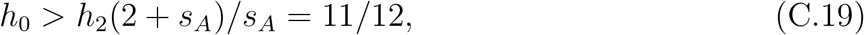

*C_AZ_* (1) is positive in spite of the local instability of the fixation state.

Key vector (B.1), which connects the local stability criterion to the Li-Price equation (20), remains valid only in the range

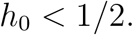

Figure B1 plots *δ*^̃^_0_ and *δ*^̃^_1_, elements of key vector (C.16), as a function of the value of *h*_0_. The relative frequency of heterozygotes (*δ*^̃^_1_) becomes non-positive for *h*_0_ *≥* 1*/*2. In addition, at *h*_0_ = 3*/*4, the principal minor Det[(***I-N***)_*n−*1_] in (B.3) passes through zero, inducing a discontinuity in the key vector (vertical line in Fig. B1).

**Figure B1:**
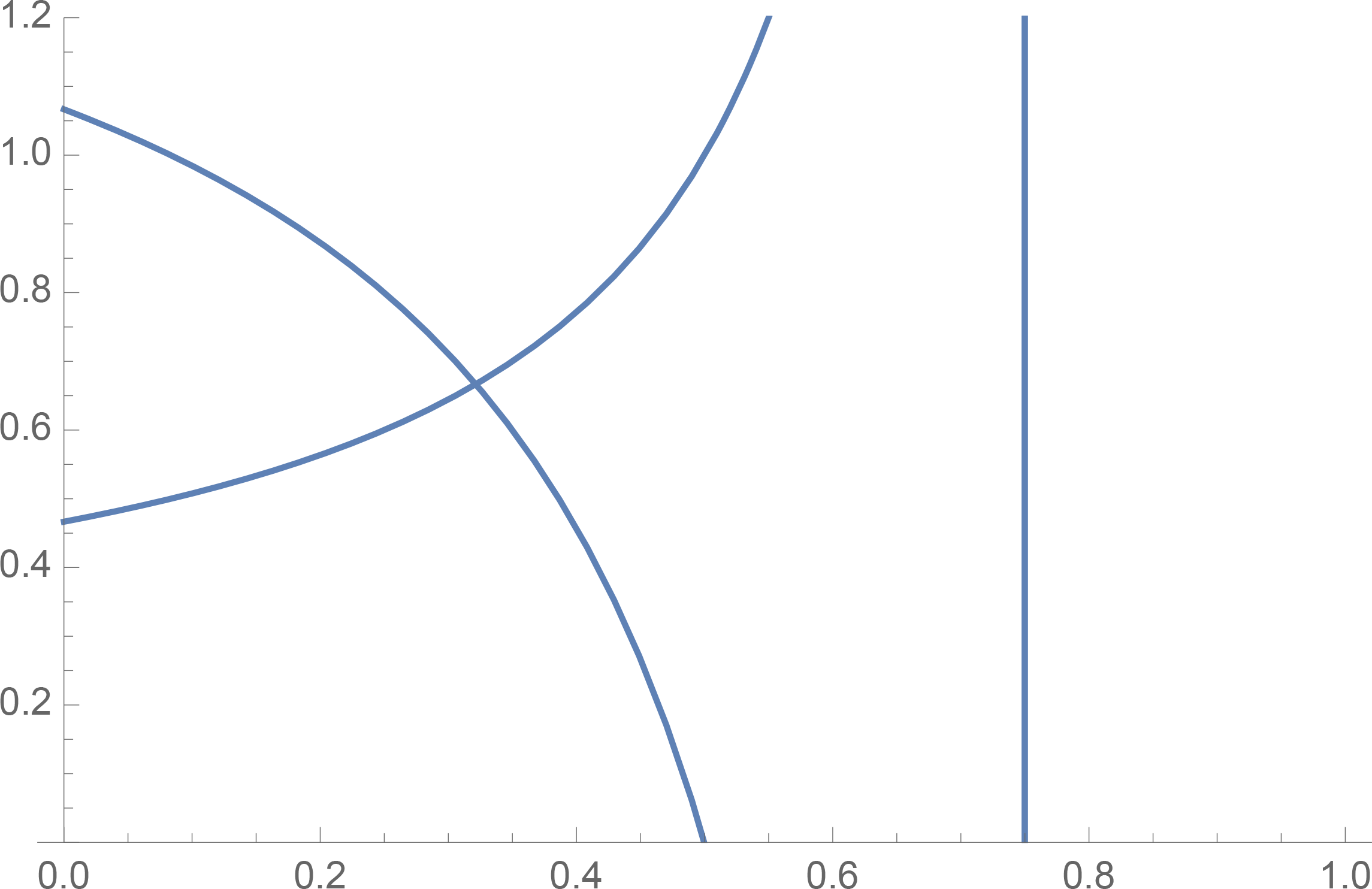
Relative magnitudes of the frequencies of rare homozygotes (*δ*^̃^_0_, increasing curve) and heterozygotes (*δ*^̃^_1_, declining curve) at the key vector (C.16) as a function of *h*_0_, the sex expression parameter associated with the rare homozygote. At the vertical bar (*h*_0_ = 3*/*4), both elements have a discontinuity, which corresponds to the passage through zero of Det[(***I** − **N***)_*n−*1_] in (B.3).

This simple example illustrates that the connection between the Li-Price equation (20) and the full local stability analysis holds only for weak selection, which corresponds under (C.17) to the range 0 *< h*_0_ *<* 1*/*2 under additivity of sex expression.

## References

Bailey, M. F. and Delph, L. F., 2007. A field guide to models of sex-ratio evolution in gynodioecious species. Oikos 116, 1609–1617.

Campbell, D. R., Weller, S. G., Sakai, A. K., Culley, T. M., Dang, P. N., and Dunbar-Wallis, A. K., 2010. Genetic variation and covariation in floral allocation of two species of *Schiedea* with contrasting levels of sexual dimorphism. Evolution 65, 757–770.

Charlesworth, B. and Charlesworth, D., 1978. A model for the evolution of dioecy and gynodioecy. Am. Nat. 112, 975–997.

Charnov, E. L., Maynard Smith, J., and Bull, J. J., 1976. Why be a hermaphrodite. Nature 263, 125–126.

Christiansen, F. B., 1991. On conditions for evolutionary stability for a continuously varying character. Am. Nat. 138, 37–50.

Collin, C. L. and Shykoff, J. A., 2003. Outcrossing rates in the gynomonoecious-gynodioecious species *Dianthus silvestris* (Caryophyllaceae). Amer. J. Bot. 90, 579–585.

Crow, J. F. and Denniston, C., 1988. Inbreeding and variance effective population numbers. Evolution 42, 482–495.

David, P., Pujol, B., Viard, F., Castella, V., and Goudet, J., 2007. Reliable selfing rate estimates from imperfect population genetic data. Mol. Ecol. 16, 2474–2487.

Denniston, C., 1978. An incorrect definition of fitness revisited. Ann. Hum. Genet. 42, 77–85.

Edwards, A. W. F., 2000. Carl Düsing (1884) on *The Regulation of the Sex-Ratio*. Theor. Pop. Biol. 58, 255–257.

Ellison, A., Cable, J., and Consuegra, S., 2011. Best of both worlds? association between outcrossing and parasite loads in a selfing fish. Evolution 65, 3021–3026.

Ellison, A., Rodriguez López, C. M., Moran, P., Breen, J., Swain, M., Megias, M., Hegary, M., Wilkinson, M., Pauluk, R., and Consuegra, S., 2016. Epigenetic regulation of sex ratios may explain natural variation in self-fertilization rates. Proc. R. Soc. Lond. B 282, 20151900.

Endler, J. A., 1980. Natural selection on color patterns in *Poecilia reticulate*. Evolution 34, 76–91.

Enjalbert, J. and David, J. L., 2000. Inferring recent outcrossing rates using multilocus individual heterozygosity: application to evolving wheat populations. Genetics 156, 1973–1982.

Eshel, I. and Motro, U., 1981. Kin selection and strong evolutionary stability of mutual help. Theor. Pop. Biol. 19, 420–433.

Ewens, W. J., 1982. On the concept of effective population size. Theor. Pop. Biol. 21, 373–378.

Fisher, R. A., 1930. The Genetical Theory of Natural Selection. Oxford Univ. Press, Oxford, first edition.

Fisher, R. A., 1941. Average excess and average effect of a gene substitution. Ann. Eugen. 11, 53–63.

Fisher, R. A., 1958. The Genetical Theory of Natural Selection. Dover, New York, second edition.

Fishman, L. and Willis, J. H., 2006. A cytonuclear incompatibility causes anther sterility in *Mimulus* hybrids. Evolution 60, 1372–1381.

Furness, A. I., Tatarenkov, A., and Avise, J. C., 2015. A genetic test for whether pairs of hermaphrodites can cross-fertilize in a selfing killifish. J. Hered. 106, 749–752.

Gantmacher, F. R., 1959. The Theory of Matrices, volume II. Chelsea Publishing Co., New York.

Kelley, J. L., Yee, M.-C., Brown, A. P., Richardson, R. R., Tatarenkov, A., Lee, C. C., Harkins, T. T., Bustamante, C. D., and Earley, R. L., 2016. The genome of the self-fertilizing mangrove rivulus fish, *Kryptolebias marmoratus*: a model for studying pheno-typic plasticity and adaptations to extreme environments. Genome Biol. Evol. 8, 2145–2154.

Laporte, V., Cuguen, J., and Couvet, D., 2000. Effective population sizes for cytoplasmic and nuclear genes in a gynodioecious species: the role of the sex determination system. Genetics 154, 447–458.

Li, C. C., 1967. Fundamental theorem of natural selection. Nature 214, 505–506.

Lloyd, D. G., 1975. The maintenance of gynodioecy and androdioecy in angiosperms. Genetica 45, 325–339.

Lloyd, D. G., 1977. Genetic and phenotypic models of natural selection. J. Theor. Biol. 69, 543–560.

Mackiewicz, M., Tatarenkov, A., Taylor, D. S., Turner, B. J., and Avise, J. C., 2006. Extensive outcrossing and androdioecy in a vertebrate species that otherwise reproduces as a self-fertilizing hermaphrodite. Proc. Natl. Acad. Sci. (USA) 103, 9924–9928.

Maynard Smith, J. and Price, G. R., 1973. The logic of animal conflict. Nature 246, 15–18.

McCauley, D. E. and Bailey, M. F., 2009. Recent advances in the study of gynodioecy: the interface between theory and empiricism. Ann. Bot. (Lond) 104, 611–620.

Price, G. R., 1970. Selection and covariance. Nature 227, 520–521.

Price, G. R., 1971. Fisher’s “fundamental theorem” made clear. Ann. Hum. Genet. 36, 126–140.

Redelings, B. D., Kumagai, S., Tatarenkov, A., Wang, L., Sakai, A. K., Weller, S. G., Culley, T. M., Avise, J. C., and Uyenoyama, M. K., 2015. A Bayesian approach to inferring rates of selfing and locus-specific mutation. Genetics 201, 1171–1188.

Reznick, D. N., Butler IV, M. J., Rodd, F. H., and Ross, P., 1996. Life-history evolution in guppies (*Poecilia reticulata*) 6. differential mortality s a mechanism for natural selection. Evolution 50, 1651–1660.

Ritland, K., 2002. Extensions of models for the estimation of mating systems using *n* independent loci. Heredity 88, 221–228.

Robertson, A., 1966. A mathematical model of the culling process in dairy cattle. Animal Production 8, 95–108.

Ross, M. D. and Weir, B. S., 1975. Maintenance of male sterility in plant populations. III. Mixed selfing and random mating. Heredity 35, 21–29.

Ross, M. D. and Weir, B. S., 1976. Maintenance of males and females in hermaphrodite populations and the evolution of dioecy. Evolution 30, 425–441.

Schultz, S. T., 1994. Nucleo-cytoplasmic male sterility and alternative routes to dioecy. Evolution 48, 1933–1945.

Serre, D., 2010. Matrices: Theory and Applications. Springer, New York.

Shaw, R. D. and Mohler, J. D., 1953. The selective significance of the sex ratio. Am. Nat. 87, 337–342.

Tatarenkov, A., Earley, R. L., Taylor, D. S., and Avise, J. C., 2012. Microevolutionary distribution of isogenicity in a self-fertilizing fish (*Kryptolebias marmoratus*) in the Florida Keys. Integr. Comp. Biol. 52, 743–752.

Taylor, P. D., 1989. Evolutionary stability in one-parameter models under weak selection. Theor. Pop. Biol. 36, 125–143.

Turner, B. J., Davis, W. P., and Taylor, D. S., 1992. Abundant males in populations of a selfing hermaphrodite fish, *Rivulus marmoratus*, from some Belize cays. J. Fish Biol. 40, 307–310.

Turner, B. J., Fisher, M. T., Taylor, D. S., Davis, W. P., and Jarrett, B. L., 2006. Evolution of ‘maleness’ and outcrossing in a population of the self-fertilizing killifish, *Kryptolebias marmoratus*. Evolutionary Ecology Research 8, 1475–1486.

Uyenoyama, M. K., 1988. On the evolution of genetic incompatibility systems. IV. Modification of response to an existing antigen polymorphism under partial selfing. Theor. Pop. Biol. 34, 347–377.

Uyenoyama, M. K., 1991. On the evolution of genetic incompatibility systems. VI. A three-locus modifier model for the origin of gametophytic self-incompatibility. Genetics 128, 453–459.

Wallace, L. E., Culley, T. M., Weller, S. G., Sakai, A. K., Kuenzi, A., Roy, T., Wagner, W. L., and Nepokroeff, M., 2011. Asymmetrical gene flow in a hybrid zone of Hawaiian Schiedea (Caryophyllaceae) species with contrasting mating systems. PLoS ONE 6, e24845. doi:10.1371/journal.pone.0024845.

Weller, S. G. and Sakai, A. K., 1991. The genetic basis of male sterility in *Schiedea* (Caryophyllaceae), an endemic Hawaiian species. Heredity 67, 265–273.

Weller, S. G. and Sakai, A. K., 2005. Inbreeding and resource allocation in *Schiedea salicaria* (Caryophyllaceae), a gynodioecious species. J. Evol. Biol. 18, 301–308.

Wolf, D. E. and Takebayashi, N., 2004. Pollen limitation and the evolution of androdioecy from dioecy. Am. Nat. 163, 122–137.

Wolff, K., Friso, B., and van Damme, J. M. M., 1988. Outcrossing rates and male sterility in natural populations of *Plantago coronopus*. Theor. Appl. Genet. 76, 190–196.

Wright, S., 1931. Evolution in Mendelian populations. Genetics 16, 97–159.

Wright, S., 1933. Inbreeding and homozygosis. Proc. Natl. Acad. Sci. (USA) 19, 411–420.

